# Computational demands shape seizure susceptibility in recurrent neural networks

**DOI:** 10.64898/2026.07.02.735135

**Authors:** Muhang Li, Sebastian Eydam, Ismaeel Ramzan, Denis Polygalov, Arthur J. Y. Huang, Ignacio Taguas, Hannah Nemeth, Dai Yanagihara, Thomas J. McHugh, Louis Kang

## Abstract

Brain areas differ in their inherent susceptibility to focal seizures, but the principles governing this risk remain unclear. While prior work has focused on anatomical and physiological factors, here we observed a fundamental contribution from the computations performed by the underlying neural network. Handcrafted and trained recurrent neural networks supporting continuous representations respond to seizure perturbations with higher activity and earlier performance decline relative to matched networks stabilizing discrete, well-separated states. Consistent with this prediction, *in vivo* recordings revealed that medial entorhinal cortex, whose grid cells exhibit continuous attractor dynamics, drives acute epileptiform discharges with stronger involvement and smoother state trajectories compared to CA3, a hippocampal subfield associated with discrete memory storage. Moreover, selective synaptic silencing demonstrated that this difference in seizure responses depends on intact entorhinal connectivity. Thus, the computations that enable neural networks to process information also influence their vulnerability to pathological transitions.

## Introduction

Seizures are states of excessive neural activity associated with neurological deficits ^1^. Focal seizures originate in a single brain area, and identifying their sites of onset and understanding their underlying pathogenesis remains an important clinical goal. While in some cases their anatomical selectivity reflects chronic pathological changes localized to one brain region, focal seizures can also arise from acute perturbations to otherwise healthy brains. Moreover, their onset can be focal even with brain-wide perturbations such as body temperature changes, drug administration, and electrolyte disturbances ^2–4^, which indicates that brain regions have inherently different susceptibilities to developing seizures. The factors that govern this intrinsic vulnerability and its variation across the brain are not fully known. Several properties involving local anatomy and physiology have been implicated in the initiation of acute seizures, including neuronal excitability, network disinhibition, and excitatory circuit motifs ^5–11^. While these properties may contribute to regional differences in seizure susceptibility, direct comparisons between brain areas and connections to broader conceptual frameworks have been lacking.

Brain areas differ not only in anatomy and physiology but also in the tasks performed by their underlying neural networks. Task computations have inherent demands on network dynamics, which may also shape pathological dynamics under ictogenic conditions. This idea is especially relevant for recurrent neural networks (RNNs), whose connectivity can amplify perturbations into seizures ^13^. RNN dynamics can be conceptualized by an energy landscape across network state space in which states with lower energy are more preferred (Fig. 1A). RNNs with a continuum of low-energy states can maintain and track continuous variables. Alternatively, RNNs with a discrete set of low-energy states can maintain and distinguish between memories, with energy barriers that prevent them from mixing. Under seizure-promoting conditions, we hypothesize that landscapes with more continuous features will provide easy dynamical paths through which excessive activity can be amplified. In contrast, the energy barriers in landscapes with more discrete features will constrain excessive activity and prevent its spread to other states represented by other neurons. The hippocampal-entorhinal system provides a well-suited biological setting for testing this idea. Its subnetworks are common initiators of focal seizures ^14^. Moreover, two of them, medial entorhinal cortex (MEC) and CA3, use recurrent connectivity to perform distinct classes of computation ^15,16^. MEC contains grid cells that encode space through continuous attractor dynamics ^17–19^, whereas CA3 is believed to store episodic memories as a discrete attractor ^20–24^.

**Figure 1.**
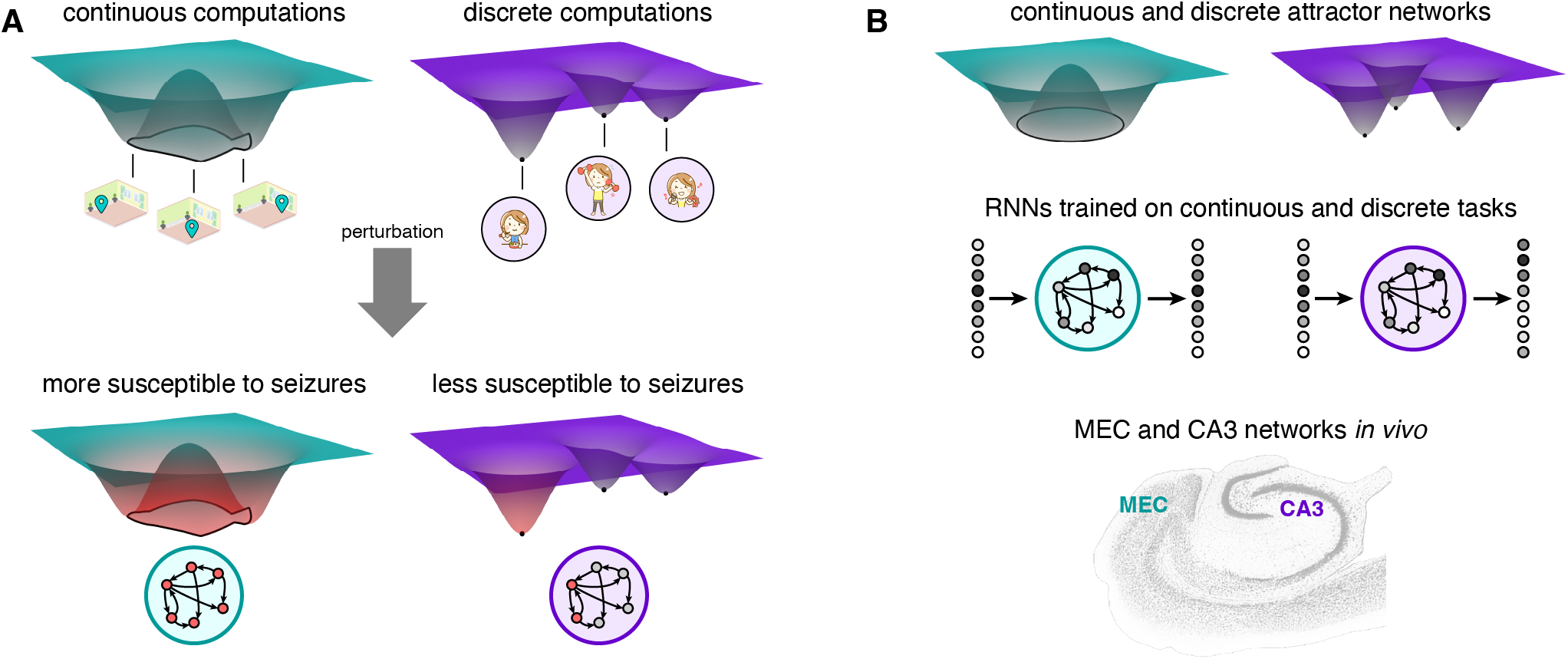
Conceptual approach. (**A**) Many networks that perform continuous memory computations, like path integration during spatial navigation, possess a continuous manifold of low-energy states. We hypothesize that excessive activity can spread along this manifold. Many networks that perform discrete memory computations, like pattern separation of episodic memories, possess high energy barriers between discrete low-energy states. We hypothesize that these barriers can constrain excessive activity. (**B**) We compare three pairs of networks with continuous and discrete counterparts: attractor networks with precise energy landscapes, RNNs trained through backpropagation, and brain areas MEC and CA3 within the mouse hippocampal-entorhinal region. Brain section image adapted from The Mouse Brain Library ^12^.

We explore seizure susceptibility in three RNN systems that perform continuous and discrete computations (Fig. 1B). First, we simulate continuous and discrete attractor networks, whose synaptic weights are designed to create precise energy landscapes. Second, we train spiking RNNs to perform either a continuous autoencoding version or a discrete classification version of the same memory task, letting optimization rather than handcrafted connectivity shape network dynamics. Third, we test these ideas experimentally through Neuropixels recordings in mouse MEC and CA3. Across these three systems, we measure population-level responses and observe internal network state dynamics as they are destabilized by acute seizure perturbations, allowing us to ask whether the computations that empower their performance also determine how they fail.

## Results

### Continuous and discrete attractor networks

We construct continuous and discrete attractor networks as periodic 2D neural sheets that contain one excitatory and one inhibitory neuron at each position (Fig. 2A; Methods). The neurons produce spikes through leaky integrate-and-fire dynamics with parameter values matched between continuous and discrete attractors. To precisely control recurrent connectivity statistics, we use binary excitatory and inhibitory synapses. Continuous and discrete attractors have the same broad inhibitory connectivity to excitatory neurons within a large distance range (Fig. 2B). The organization of excitatory outputs differentiates the two network topologies. The continuous attractor has local excitatory connectivity to other excitatory and inhibitory neurons within a short distance. In contrast, excitatory neurons in the discrete attractor are partitioned non-locally into disjoint assemblies of evenly spaced neurons throughout the neural sheet. Each excitatory neuron connects to a subset of other excitatory neurons within the same assembly and to the corresponding inhibitory neurons at the same positions. Importantly, we tune the continuous connectivity distance and the connectivity density within discrete assemblies such that each neuron in both continuous and discrete attractor networks has the same number of incoming and outgoing synapses.

**Figure 2.**
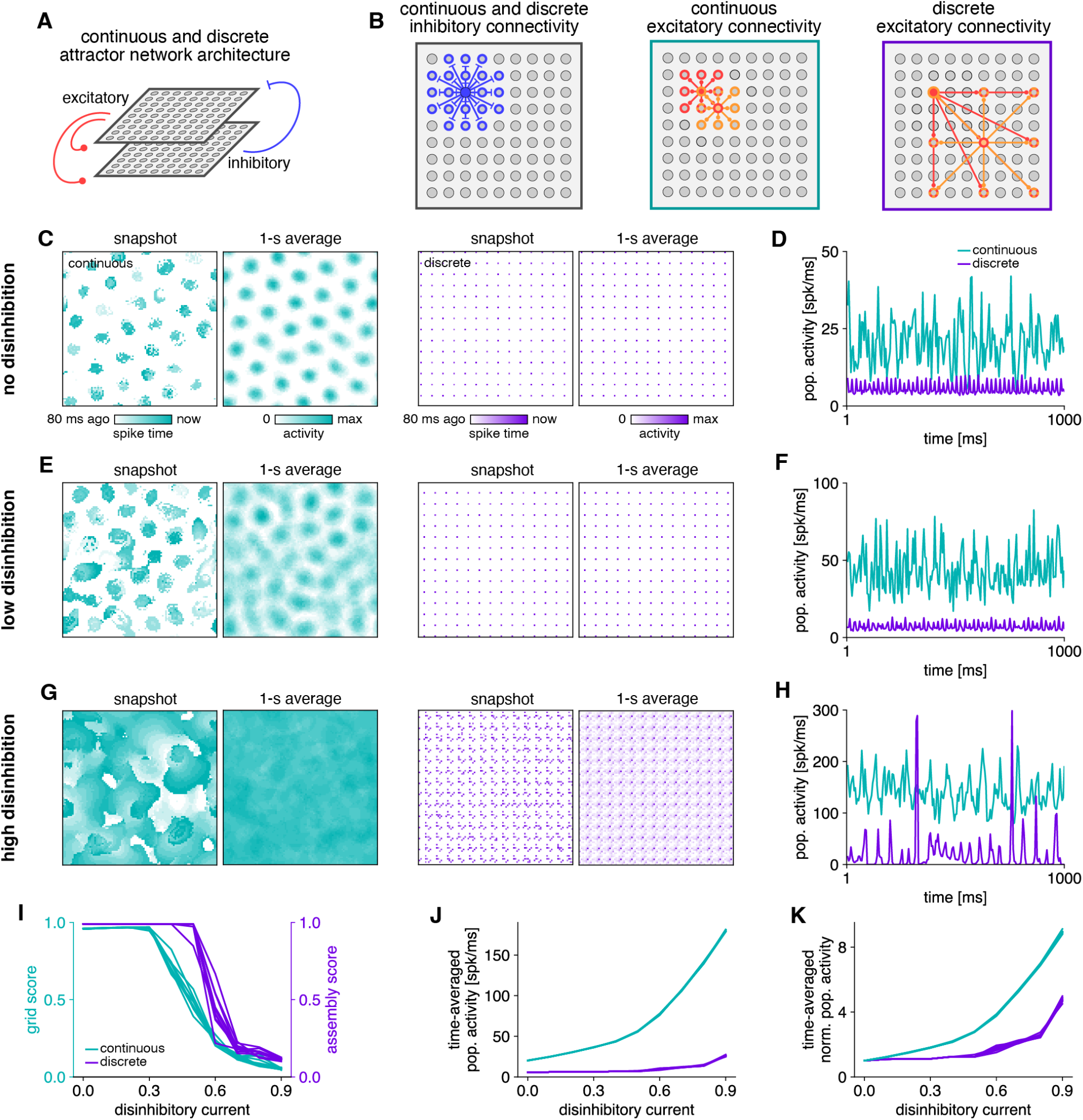
Continuous and discrete attractor networks with negative current applied to the inhibitory population. (**A**) Spiking neurons are arranged in a 2D neural sheet with overlapping excitatory and inhibitory populations and periodic boundaries. Recurrent connections exist within and across populations, except for inhibitory-to-inhibitory connections. (**B**) Inhibitory neurons have broad local connectivity in both continuous and discrete attractor networks. In continuous attractors, excitatory neurons have narrow local connectivity similar to grid cell attractor networks. In discrete attractors, excitatory neurons have long-range connectivity to other neurons within the same assembly, which are regularly spaced throughout the neural sheet. Example neurons and their synaptic targets are highlighted with filled and open colored circles, respectively. (**C,D**) Network activity at baseline. Both continuous and discrete attractor networks can maintain stable attractor states. (**C**) Excitatory neuron activity. Each pixel corresponds to one neuron. (**D**) Excitatory population activity averaged over 5-ms time bins. (**E,F**) Similar to **C,D**, but for low disinhibitory current 0.5. Activity bumps expand and merge in the continuous attractor while the discrete attractor preserves its operation. (**G,H**) Similar to **C,D**, but for high disinhibitory current 0.9. The continuous attractor exhibits traveling wavefronts with high persistent activity. The discrete attractor simultaneously activates multiple assemblies in population activity bursts. (**I**) Both networks lose attractor function under increasing disinhibitory current, and continuous attractors degrade earlier. (**J, K**) Continuous attractors maintain higher activity levels in response to increasing disinhibitory current. In **K**, the activity of each network in **J** is divided by its value at disinhibitory current 0. **I**–**K** show data from 10 replicate simulations. Each line corresponds to one replica.

These networks indeed function as continuous and discrete attractors. Continuous attractors with similar architectures have been used to model entorhinal grid cells ^25,26^, and accordingly, our network maintains continuous attractor states consisting of activity bumps that form a triangular grid. Figure 2C (left) shows one such state, and the complete 2D attractor manifold can be traversed through uniform translations of the bumps. Our discrete attractor network activates its neural assemblies as discrete attractor states. Figure 2C (right) shows one such assembly, and every other similarly spaced square sublattice represents another attractor state.

To these functional networks, we apply a disinhibitory perturbation with a current that effectively lowers the resting membrane potential of the inhibitory neurons. With a small perturbation, the continuous attractor no longer maintains a stable grid as its bumps grow, drift, and intermittently merge (Fig. 2E, left; Supp. Fig. 1). This corresponds to an increase in population activity relative to that of the discrete attractor (Fig. 2D,F), which still maintains stable attractor states (Fig. 2E, right). Under high disinhibition, the continuous attractor produces activity wavefronts that continually propagate across the neural sheet (Fig. 2G, left; Supp. Fig. 1). Neurons participate sequentially in these wavefronts, which permits a high level of population activity despite each neuron’s relative refractory period after spiking and resetting its membrane potential (Fig. 2H). Meanwhile, the discrete attractor mixes multiple assemblies (Fig. 2G, right). The dense recurrent excitation within each assembly causes neurons to spike and become relatively refractory in synchrony, so the network exhibits population bursts instead of sustaining high population activity (Fig. 2H).

Overall, disinhibitory current disrupts both networks’ ability to maintain their original attractor states, as quantified by grid and assembly scores (Fig. 2I; Methods). Continuous attractors experience earlier performance degradation, indicating that their computations, which require a higher degree of symmetry, are more unstable. They also respond with higher population activity (Fig. 2J,K). The greater vulnerability of continuous attractor networks persists over a wide range of network parameters and largely holds across alternative perturbations involving synaptic strength (Supp. Fig. 2).

### RNNs trained on continuous and discrete tasks

To explore the generality of our observations beyond attractor networks with handcrafted weights, we turn to trained RNNs, whose connectivities are defined through task optimization. Our RNNs have a recurrent layer of spiking leaky integrate-and-fire neurons, which receive inputs from a layer of rate-based neurons and send outputs to another layer of rate-based neurons of equal width (Fig. 3A; Methods). The input and output layers are linearly ordered with periodic boundaries. The RNNs are trained to perform one of two memory tasks (Fig. 3B). In both tasks, the input layer presents a bump of activity centered around a particular position for 20 time steps. In the continuous memory task, the RNN must produce an activity bump at the same position in the output layer for time steps 21 to 100. In the discrete memory task, the RNN must determine whether the input bump is centered in the top or bottom half of the input layer, and then it must produce an activity bump centered within the correct half of the output layer for time steps 21 to 100. Thus, the location of the output bump is the only difference between the two tasks; it is determined via a continuous or a discrete input–output mapping.

**Figure 3.**
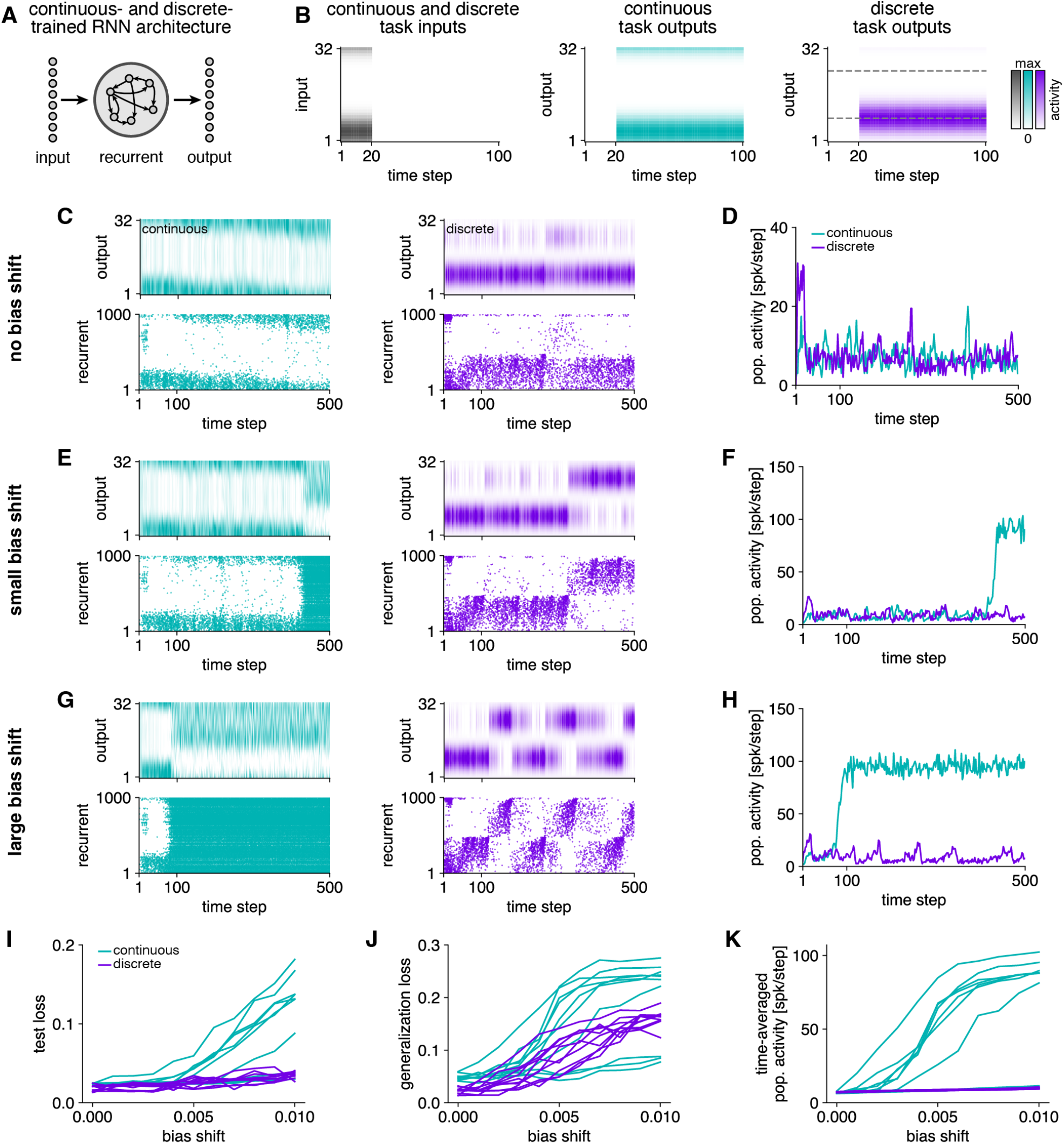
Continuous- and discrete-trained RNNs with a positive shift in recurrent layer biases. (**A**) A recurrent layer of spiking neurons receives inputs from and sends outputs to layers of rate-based neurons arranged linearly with periodic boundaries. (**B**) The input is a transient activity bump in both continuous and discrete tasks. In the continuous task, the RNN must output a bump at the same location. In the discrete task, the RNN must output a bump at one of two locations depending on the input bump location. (**C,D**) Trained RNN activity at baseline. Both continuous- and discrete-trained networks can maintain an output bump in the correct position and partially generalize beyond the 100 training time steps. (**C**) Outputs and recurrent layer spike rasters. (**D**) Population activity averaged over 2-step time bins. (**E,F**) Similar to **C,D**, but for low bias shift 0.004. The continuous-trained RNN becomes unstable and maintains high persistent activity. The discrete-trained RNN transitions to the incorrect target bump. (**G,H**) Similar to **C,D**, but for high bias shift 0.008. The continuous-trained RNN becomes unstable before 100 time steps and maintains high persistent activity. The discrete-trained RNN maintains stability within 100 time steps but thereafter fluctuates between target bumps. (**I,J**) Continuous-trained RNNs show substantially higher test loss under increasing bias shift, and both networks show higher generalization loss. (**K**) Continuous-trained RNNs maintain higher activity levels in response to increasing bias shift. **I**–**K** show data from 10 replicate trained RNNs tested on 32 samples each. Each line corresponds to one replica averaged over test samples.

The RNNs are trained end-to-end with backpropagation through time and surrogate gradients (Methods; Supp. Fig. 3). The total loss equals a mean-squared-error task loss plus regularizer terms that encourage the continuous- and discrete-trained RNNs to solve their tasks using the same level of recurrent activity and distribution of recurrent weights and biases. With these regularizers, differences between the networks observed during perturbations can be more confidently ascribed to task identity rather than differences in incidental network properties. The trained RNNs can indeed perform their tasks and maintain output bumps in roughly the correct location even beyond 100 time steps, demonstrating temporal generalization (Fig. 3C). Sorted spike raster plots illustrate that the recurrent layer state tracks the output bump. Our regularizer successfully matches the activity levels of continuous- and discrete-trained RNNs (Fig. 3D).

To model an acute ictogenic perturbation, we freeze all weights and biases after training and increase all recurrent biases by a small positive value. With a low bias shift, the RNNs can still operate over the 100 time steps of training, but they break down sometime thereafter (Fig. 3E,F). The continuous-trained RNN transitions to a sustained high-activity regime that no longer outputs a bump. The discrete-trained RNN maintains a compact bump without widespread activity, but it intermittently jumps from the correct target location to the other one. A high bias shift exacerbates these instabilities, with the continuous-trained RNN transitioning quickly to the high-activity regime and the discrete-trained network jumping frequently between its two target bumps (Fig. 3G,H).

Overall, shifting recurrent biases decreases RNN performance. The continuous-trained RNNs start to fail even within the training duration of 100 time steps, whereas the discrete-trained RNNs are more resilient (Fig. 3I). Beyond 100 time steps, both networks show impaired generalization (Fig. 3J). Notably, the two networks lose task performance through distinct dynamical regimes. The continuous-trained RNNs sustain high activity spread across states present in the unperturbed condition, whereas the activity in discrete-trained RNNs is largely limited to their unperturbed states (Fig. 3K). These observations of higher perturbed activity and more fragile task performance in the continuous-trained RNN are largely maintained across a wide range of network hyperparameters and for an alternative perturbation of recurrent weights (Supp. Figs. 4 and 5). Analysis of recurrent weight matrices indicates that these networks have learned attractor connectivity structures broadly at the network scale, though with substantial synaptic variability at the neuron scale (Supp. Figs. 3 and 5).

The close agreement across attractor networks and trained RNNs, despite their architectural and conceptual differences, reinforces the connection between computational demands under normal regimes and seizure-like dynamics under perturbed regimes. In both systems, the continuous networks are more strongly affected by imbalances of excitation over inhibition, as manifested by loss of normal functioning and excessive neural activity.

### High-density neural recordings in MEC and CA3

To test whether our simulation results apply to biological neural networks, we recorded simultaneously from mouse MEC layers II and III (L2/3) and CA3 using Neuropixels probes (Fig. 4A; Supp. Fig. 6; Methods). In our simulations, we applied an acute, uniform perturbation to both continuous and discrete networks for a fair comparison between them. We experimentally implemented this manipulation with an acute intraperitoneal (i.p.) injection of pentylenetetrazole (PTZ), a GABA_A_ receptor antagonist ^27,28^. To study the intrinsic susceptibilities of these brain regions and reduce confounding from external input or behavior, we recorded under urethane anesthesia, a state which preserves physiological activity in the hippocampal-entorhinal circuit ^29–31^. Upon a high-dose injection of PTZ, epileptiform discharges were visible in the local field potential (LFP) of both MEC and CA3 in wildtype (wt) mice (Fig. 4B). To examine the differential contributions of MEC and CA3 to this pathophysiology, we virally expressed the tetanus toxin light chain gene (tetX) specifically in superficial MEC excitatory neurons (tetMEC) or in CA3 pyramidal cells (tetCA3) in separate animal cohorts (Fig. 4C,E; Supp. Figs. 6 and 7; Methods). Tetanus toxin prevents neurotransmitter release at axon terminals, which blocks synaptic transmission without impairing action potential generation. Under PTZ, tetMEC mice exhibited markedly weaker discharges in both MEC and CA3 (Fig. 4D), while in tetCA3 mice, discharges decreased in CA3 but persisted in MEC (Fig. 4F; Supp. Fig. 8). Quantification confirmed that silencing MEC synaptic outputs significantly decreased peak LFP amplitude in both MEC and CA3 compared to wt mice, demonstrating their necessity for discharges in both regions (Fig. 4G–I). While silencing CA3 synaptic outputs significantly decreased CA3 peak amplitudes, MEC peak amplitudes remained similar to those of wt mice, suggesting a more limited role for CA3. Similar results were observed for LFP power across multiple frequency bands (Supp. Fig. 9).

**Figure 4.**
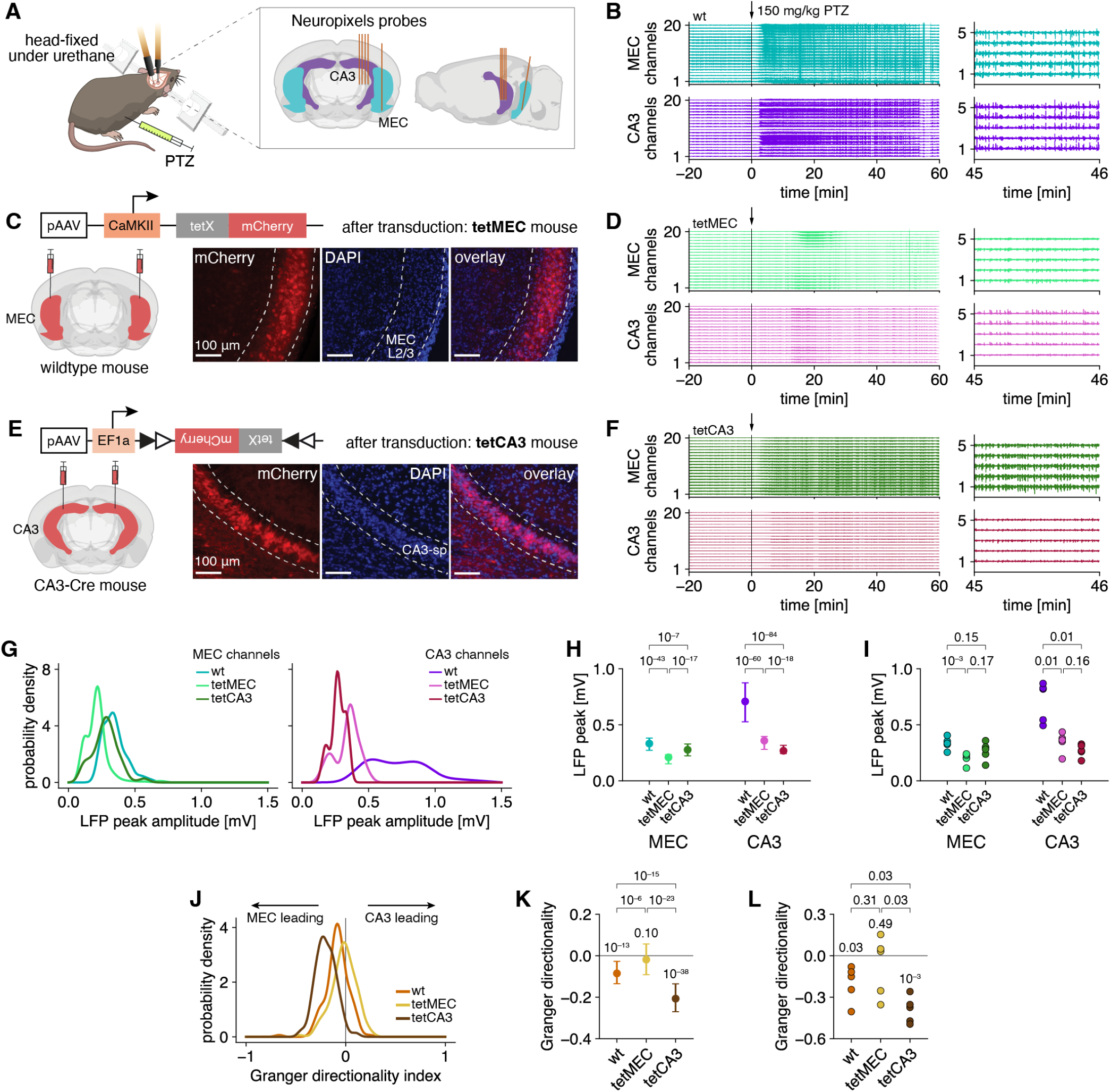
MEC and CA3 LFP analysis under high-dose PTZ. (**A**) We simultaneously record from MEC and CA3 in anesthetized mice using Neuropixels probes. (**B**) MEC and CA3 LFP signals in a wt mouse. High-dose PTZ is injected at 0 min. (**C**) Outputs of superficial MEC excitatory neurons are silenced by selective expression of tetanus toxin (tetMEC) with mCherry marker. (**D**) Similar to **B**, but for a tetMEC mouse. (**E**) Outputs of CA3 pyramidal cells are silenced by selective expression of tetanus toxin (tetCA3) with mCherry marker. sp: stratum pyramidale. (**F**) Similar to **B**, but for a tetCA3 mouse. (**G**–**I**) PTZ-induced epileptiform discharges are suppressed in the MEC and CA3 of tetMEC mice and suppressed only in the CA3 of tetCA3 mice. (**G**) Kernel densities of LFP peak amplitudes per channel, defined as the 95th percentile Hilbert amplitude between 30–60 min, aggregated across animals. (**H**) Medians and interquartile ranges of the data in **G.** (**I**) Medians of LFP peak amplitudes per animal. (**J**–**L**) MEC LFP precedes CA3 LFP in wt mice during PTZ seizures. This relationship is weakened in tetMEC mice and strengthened in tetCA3 mice. (**J**) Kernel densities of the Granger directionality index per 1-min LFP window between 30–60 min aggregated across animals. Negative and positive values correspond to MEC and CA3 leading, respectively. (**K**) Medians and interquartile ranges of the data in **J**. (**L**) Granger directionality index between 30–60 min per animal. **G**–**L** show data for 5 wt, 5 tetMEC, and 5 tetCA3 animals. In **H, K**, numbers above brackets indicate *p*-values for Mood’s tests comparing medians. In **I, L**, numbers above brackets indicate *p*-values for unpaired two-sample *t*-tests comparing means. In **K**, numbers below brackets indicate *p*-values for one-sample quantile tests comparing the median with 0. In **L**, numbers below brackets indicate *p*-values for one-sample *t*-tests comparing the mean with 0.

If MEC activity were to drive discharges in CA3, we would expect LFP signals in MEC to lead those of CA3. This temporal relationship was quantified by the Granger directionality index between MEC and CA3, which indicated that MEC leading CA3 is the dominant direction of temporal precedence for wt mice in the seizure regime (Fig. 4J–L; Methods). As expected, silencing MEC outputs shifted the directionality towards CA3, and silencing CA3 outputs shifted it towards MEC. These results support our prediction that the MEC recurrent network, which performs continuous computations, is more influential during ictogenesis compared to the CA3 recurrent network, which performs discrete computations.

Next, we investigate the network mechanisms of PTZ-induced destabilization. Our simulations predict that continuous and discrete networks should exhibit distinct responses to mild perturbations that destabilize attractor states without causing seizures. Continuous attractors should drift along the low-dimensional attractor manifold with high network state overlap from one moment to the next, while discrete attractors should hop among attractor states spread out in high-dimensional state space with low temporal overlap (Fig. 5A). These expected dynamics are confirmed by simulations (Fig. 5B) and quantified by *temporal continuity*, defined as the cosine similarity of the detrended network state between successive time bins (Methods). Continuous attractors exhibit higher temporal continuity than discrete attractors, which have values closer to zero (Fig. 5C,D).

**Figure 5.**
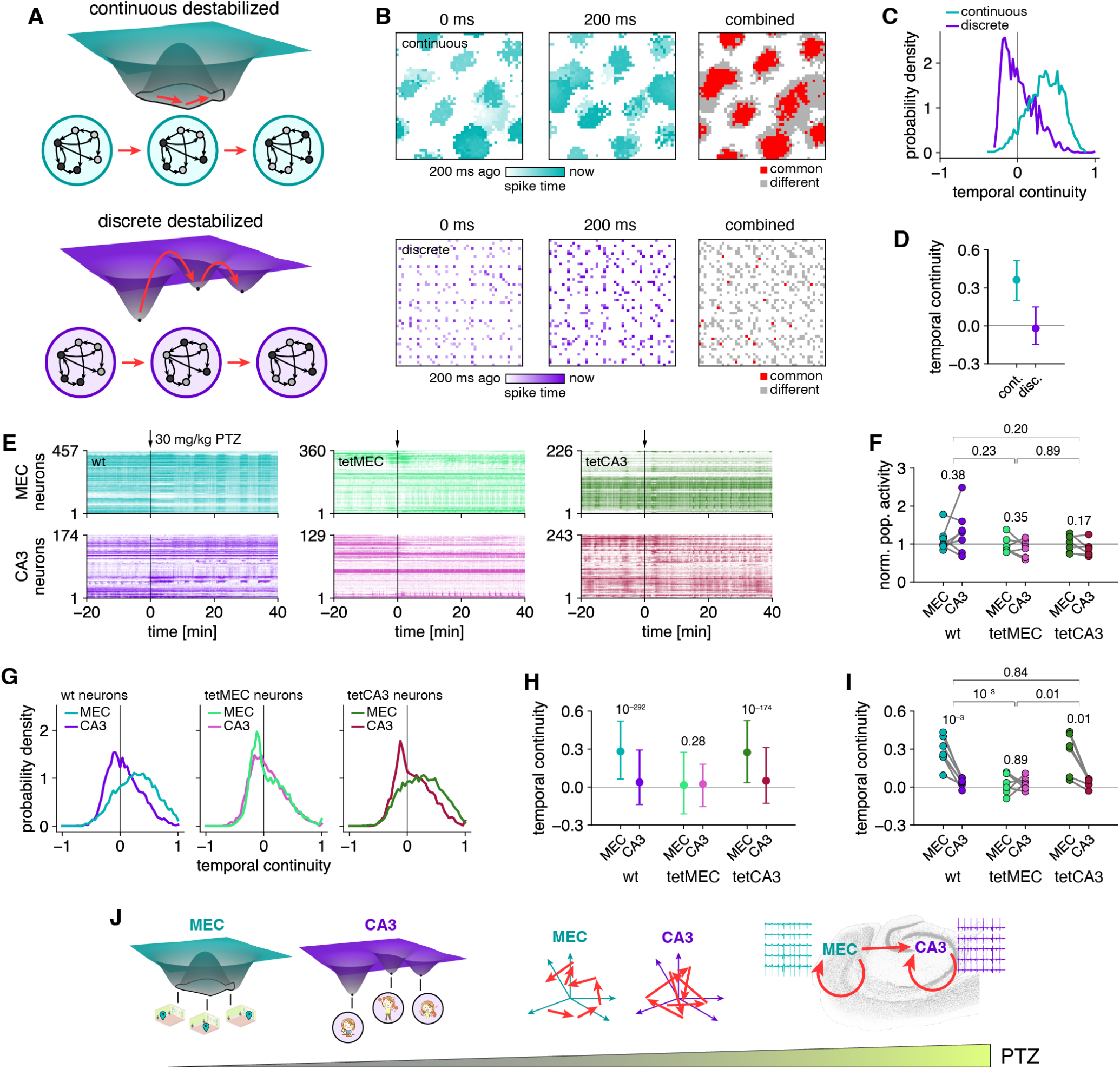
MEC and CA3 single-unit analysis under low-dose PTZ. (**A**) Under a weak perturbation that destabilizes attractor states without causing seizures, we expect the network state to change smoothly with high temporal overlap in continuous networks and abruptly with low temporal overlap in discrete networks. (**B**–**D**) Example attractor network simulations under destabilizing conditions. (**B**) Excitatory neuron activity during consecutive 200-ms time bins (left, middle) and neurons active in one or both bins (right). (**C**) Histograms of temporal continuity, which is the cosine similarity between detrended network states in successive 200-ms time bins, for a continuous and a discrete attractor simulation. (**D**) Medians and interquartile ranges of the data in **C.**(**E**) MEC and CA3 spike rasters in a wt (left), a tetMEC (middle), and a tetCA3 (right) mouse. Low-dose PTZ is injected at 0 min. (**F**) Average population firing rates between 10–40 min per animal normalized by their values before PTZ injection. (**G**–**I**) MEC and CA3 network states in wt mice exhibit high and low temporal continuity, respectively, under a weak PTZ perturbation. This effect is disrupted in tetMEC mice but preserved in tetCA3 mice. (**G**) Histograms of temporal continuity between 10–40 min aggregated across animals. (**H**) Medians and interquartile ranges of the data in **G**. (**I**) Medians of temporal continuity between 10–40 min per animal. (**J**) Conceptual model for our results. Because MEC and CA3 perform different computations under normal conditions, their network states are destabilized through distinct dynamics by small PTZ perturbations, and they make unequal contributions to seizure discharges under large PTZ perturbations. Brain section image adapted from The Mouse Brain Library ^12^. **F**–**I** show data from 6 wt, 6 tetMEC, and 6 tetCA3 mice. In **F, I**, numbers above brackets indicate *p*-values for unpaired two-sample *t*-tests comparing means of differences between paired MEC and CA3 data. Numbers below brackets indicate *p*-values for paired two-sample *t*-tests comparing means. In **H**, numbers indicate *p*-values for Mood’s tests comparing medians.

To experimentally test these predictions, we used the same recording paradigm, except with a low PTZ dose to perturb the network without entering the seizure regime. Without large epileptiform discharges that can overshadow action potential waveforms, we could perform spike sorting to yield simultaneously recorded neurons in MEC and CA3 (Fig. 5E; Supp. Fig. 10; Methods). These regions showed no significant differences in their overall levels of population activity (Fig. 5F). However, MEC and CA3 network states in wt mice demonstrated markedly different distributions of temporal continuity that agree with those of continuous and discrete attractor networks, respectively (Fig. 5G–I; Supp. Fig. 11). Moreover, in tetMEC mice, MEC temporal continuity decreased towards zero, demonstrating that the overlapping trajectories traversed under PTZ can only be accessed with MEC synaptic outputs intact. Silencing CA3 synaptic outputs in tetCA3 mice did not affect MEC temporal continuity. This agreement between experiment and simulation suggests that the MEC response to PTZ administration reflects continuous features in its network dynamics that are established through recurrent connectivity.

In summary, our experimental results support an explanatory model in which an ictogenic perturbation, in the form of PTZ administration, first induces MEC population activity to drift through nearby states while the CA3 network state fluctuates with less temporal continuity (Fig. 5J). MEC synaptic outputs are required for its overlapping trajectories, independently of CA3. With a large enough perturbation, MEC becomes fully destabilized and exhibits large seizure discharges facilitated by its synaptic outputs, which also provide a necessary contribution to large seizure discharges in CA3.

## Discussion

Our results establish a link between network computation and seizure susceptibility through three complementary approaches that impose computational demands in distinct ways. The demands of attractor networks are structurally defined by choosing their synaptic connectivities, those of trained RNNs are functionally defined by the tasks for which they are optimized, and those of networks in the hippocampal-entorhinal region arise through biological processes. In all three cases, we observed that the network performing more continuous computations responds more intensely to ictogenic perturbations than its discrete counterpart. Continuous attractors and continuous-trained RNNs maintain higher activity levels and demonstrate earlier task degradation across wide ranges of model configurations. *In vivo*, MEC synaptic outputs drive the amplification of epileptiform discharges in both MEC and CA3 with destabilization dynamics characteristic of our attractor simulations.

Our findings interface with previous observations related to seizures. Many reports have also identified entorhinal cortex as a driver of seizures ^32–36^. Meanwhile, CA3 has been observed to generate interictal spikes, which are brief bursts of electrical activity unaccompanied by obvious neurological impairments ^33,37–40^. This phenomenon resembles dynamics in our discrete attractors and discrete-trained RNNs—they demonstrate synchronized bursts of activity, and the latter can even maintain some degree of task performance throughout these bursts (Figs. 2H, 3E, and 3F; Supp. Figs. 4 and 5). Our simulations suggest that the discrete topology of CA3 yields interictal spikes under seizure-promoting perturbations, in contrast to the continuous topology that drives ictal discharges in MEC. Moreover, seizures in our continuous attractors typically manifest as traveling wavefronts of neural activity (Fig. 2G; Supp. Figs. 1 and 2), which have been previously observed in animal experiments, patient recordings, and simulations ^10,41^.

Although MEC and CA3 contain heterogeneous neural populations and exhibit complex dynamics that deviate from idealized attractor models, our comparison between them is best understood as a contrast between networks skewed towards different computational topologies. Moreover, focal seizures arise within many other brain regions, and further experimental efforts can determine how broadly our ideas apply to contrasts in computation throughout the brain. Future theoretical extensions include the implementation of richer neuronal dynamics and interactions across brain regions ^7,42,43^. Still, our networks can clearly illustrate our conceptual ideas, and they can demonstrate excessive activity and functional impairment, which are defining features of seizures common to all types ^1^.

Our work reflects broader connections between biological function and dysfunction. Computational psychiatry uses cognitive computations and their alterations to address mental disorders ^44–46^; likewise, our results suggest that neural network computations may provide insight into neurological disorders. More generally, the function–dysfunction duality we identify is conceptually related to the performance–robustness trade-off in control theory ^47^, in which more responsive feedback systems are often less stable, and to the fluctuation–dissipation theorem in statistical physics ^48^, in which thermodynamic systems that respond more strongly to external inputs also exhibit larger internal fluctuations. Here, we have shown how neural systems that can responsively encode continuous changes, in contrast to settling into rigid, discrete states, inherently suffer from greater seizure vulnerability.

## Methods

### Attractor networks

#### Network architecture and dynamics

Our attractor networks consist of a 2D neural sheet with periodic boundary conditions that contains overlapping excitatory and inhibitory populations *P ∈ {*E, I*}*. One excitatory neuron and one inhibitory neuron are located at each position **r** = (*i, j*) *∈ {*1, …, *N*} ×{1, …, *N*}. Each neuron is described by a membrane potential *v*_*P*_ (**r**, *t*) and a spike indicator *s*_*P*_ (**r**, *t*). If the potential exceeds a threshold of 1 at time *t*, the neuron fires a spike and its potential is reset to 0: *s*_*P*_ (**r**, *t*) *←* 1 and *v*_*P*_ (**r**, *t*) *←* 0. Otherwise, *s*_*P*_ (**r**, *t*) = 0. We prevented *v*_*P*_ from decreasing below −2 to limit hyperpolarization.

The membrane potential evolves according to leaky integrator dynamics:

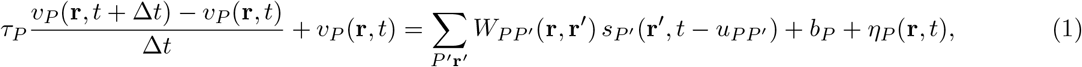

where 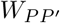is the recurrent connectivity from population *P* ^*′*^ to *P, b*_*P*_ is a constant membrane current that sets the resting membrane potential, *η*_*P*_ represents independent, zero-mean Gaussian noise, *τ*_*P*_ is the membrane time constant, and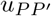is the synaptic delay. We typically set *b*_E_ *>* 1 so excitatory neurons are intrinsically driven to fire, and we set *b*_I_ *<* 1 so inhibitory neurons will not fire unless they receive synaptic input from excitatory neurons.

#### Continuous and discrete connectivity

Our networks contain I-to-E, E-to-E, and E-to-I connections with binary synapses whose strengths are constant in each of these three directions. There are no I-to-I connections: *W*_II_ = 0.

Both continuous and discrete attractor networks share the same I-to-E connectivity based on distance along the periodic neural sheet. Using the 1D periodic distance

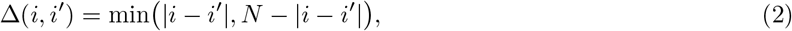

the 2D neural sheet distance becomes

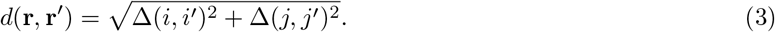

Inhibitory neurons connect to excitatory neurons located between 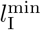and 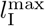away:

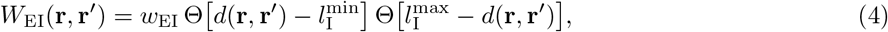

where Θ is the Heaviside step function and *w*_EI_ is the I-to-E synaptic strength.

For continuous attractor networks, excitatory neurons connect to other excitatory neurons and inhibitory neurons located within 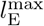away:

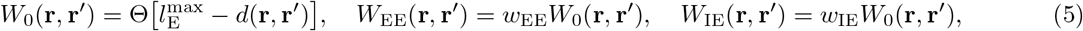

where *w*_EE_ is the E-to-E synaptic strength and *w*_IE_ is the E-to-I synaptic strength. This connectivity profile yields activity bumps that form a triangular grid on the neural sheet.

For discrete attractor networks, connectivity is controlled by setting *q* and *g*, which are two integers that divide *N* and *g* must be odd. We first partitioned the neural sheet into *q*^2^ square sublattices with spacing *q*. Excitatory neurons send connections to other excitatory and inhibitory neurons within the same sublattice as follows. We connected each position to *g*^2^ neighbors within the same sublattice:

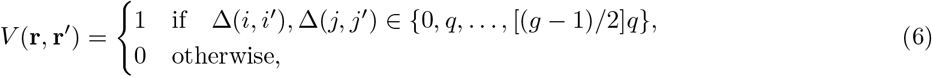

with Δ defined in Eq. 2. We then applied a stochastic degree-preserving edge swapping algorithm within each sublattice to distribute its connections non-locally without changing the number *g*^2^ of incoming and outgoing connections at each position ^49^. We scaled the output of this synaptic permutation to obtain the E-to-E and E-to-I connectivity:

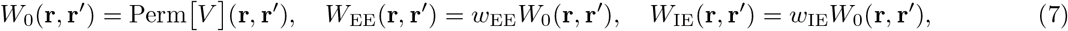

where *w*_EE_ is the E-to-E synaptic strength and *w*_IE_ is the E-to-I synaptic strength. The abundant E-to-E connections within sublattices and their absence across sublattices cause each sublattice to function as a neural assembly. With this construction method, we could control the number of assemblies *q*^2^ and the number of incoming and outgoing excitatory synapses per neuron *g*^2^, while minimizing structured connectivity patterns within assemblies that may lead to stereotyped dynamics.

#### Grid and assembly scores

The grid score measures the functional ability of the continuous attractor network to maintain an attractor state, which manifests as a triangular lattice of neural activity bumps in our case. To detect this lattice, we collected spike counts *c*(**r**) within a 200-ms time window and computed 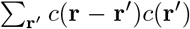, which is then normalized to yield the 2D autocorrelation function *C*(**r**). Following previous reports ^50^, we integrated *C*(**r**) radially between its first two radial minima, a region that contains its first peaks beyond the one at the origin, to obtain the angular autocorrelation *C*(*θ*) as a function of polar angle *θ*. We calculated its discrete Fourier transform: 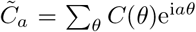 for component indices *a*. The grid score is defined as the fraction of oscillatory power in the 6th component of 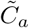:

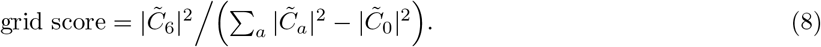

We typically present its average over the simulation.

The assembly score measures the functional ability of the discrete attractor network to maintain an attractor state, which manifests as activity concentrated in one neural assembly in our case. To compute it, we collected spike counts *c*(**r**) within a 200-ms time window and summed across neurons within each assembly sublattice: 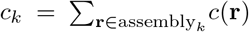for assemblies *k{* 1, …, *q*^2^*}*. We also summed across all neurons:*c* = ∑_r_ *c*(**r**). The assembly score is defined to be the normalized Herfindahl-Hirschman index, which here measures the concentration of spikes within a single assembly:

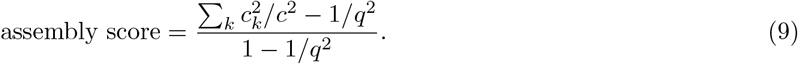

We typically present its average over the simulation.

#### Simulation protocols

We initialized attractor networks with membrane potentials consistent with an attractor state. For the continuous attractor network, we first identified the steady-state triangular lattice approached by the network over a long simulation with random initial potentials. We generated a membrane potential profile by placing 2D Gaussian bumps at each lattice point with radius 0.1 times the lattice spacing. The profile was rescaled to span membrane potential values from 0.5 to 0.95 and was used to initialize the continuous attractor networks. For the discrete attractors, we randomly chose one of the assemblies and initialized its neurons with uniformly random membrane potentials between 0.5 and 0.95; all other neurons were given potentials 0. We finally added a small amount of Gaussian noise to each neuron in both networks. In the discrete attractor network, the connectivity matrix was also generated through a stochastic permutation process, so a different realization was used in each replica.

Unless otherwise specified, we ran each simulation for 20 s and discarded data from the first 10 s. Standard unperturbed network parameters are provided in Table 1. Each perturbation is a change in these network parameters and was applied from the start of the simulation. Disinhibitory current is defined as the decrease in inhibitory current *b*_I_ from the unperturbed value. E-to-I weight decrease and I-to-E weight increase are defined to be changes in weights *w*_IE_ and *w*_EI_, respectively, from the unperturbed value.

**Table 1.** Standard parameters for continuous and discrete attractor networks unless otherwise noted.

| Parameter | Variable | Value |
| --- | --- | --- |
| number of neurons per population | $N^2$ | $98^2$ |
| time step | $\Delta t$ | 1 ms |
| inh. membrane time constant | $\tau_I$ | 20 ms |
| exc. membrane time constant | $\tau_E$ | 40 ms |
| inh. membrane current | $b_I$ | 0.6 |
| exc. membrane current | $b_E$ | 1.05 |
| inh. noise magnitude | $\text{std}[\eta_I]$ | 0.05 |
| exc. noise magnitude | $\text{std}[\eta_E]$ | 0.02 |
| I-to-E synaptic strength | $w_{EI}$ | -0.65 |
| E-to-E synaptic strength | $w_{EE}$ | 0.65 |
| E-to-I synaptic strength | $w_{IE}$ | 0.65 |
| I-to-E synaptic delay | $u_{EI}$ | 2 ms |
| E-to-E synaptic delay | $u_{EE}$ | 5 ms |
| E-to-I synaptic delay | $u_{IE}$ | 2 ms |
| minimum inh. connectivity range | $l_I^{\min}$ | 2 |
| maximum inh. connectivity range | $l_I^{\max}$ | 12 |
| (continuous only) maximum exc. connectivity range | $l_E^{\max}$ | 4 |
| (discrete only) number of assemblies | $q^2$ | $14^2$ |
| (discrete only) number of exc. synapses per neuron | $g^2$ | $7^2$ |

For the simulations in Fig. 5B–D used for temporal continuity analysis, we ran each simulation for 123 s and discarded data from the first 3 s. Differences from Table 1 are *N* ^2^ = 49^2^, *b*_E_ = 1.0, *b*_I_ = 0.3, std[*η*_E_] = 0.05, *w*_EI_ =−2, *w*_EE_ = *w*_IE_ = 0.5.

For the simulations in Supp. Fig. 2F,G comprising the parameter grid search, we ran each simulation for 11 s and discarded data from the first 1 s. We tested all 240 combinations of these parameters that differ from Table 1: *b*_E_ *∈* {1.02, 1.04, 1.06}, *w*_EI_ *∈* {−0.8, −0.7, −0.6, −0.5*}, w*_EE_ *∈* {0.6, 0.7, 0.8, 0.9}, and *w*_IE_ *∈* {0.4, 0.5, 0.6, 0.7, 0.8}. For each combination, we ran one continuous and one discrete attractor simulation for each value of *b*_I_ *∈* {0.8, 0.7, …, −0.4}. We searched for parameter combinations that could model a seizure-like transition under a disinhibitory current perturbation by satisfying two requirements. First, the combination must capture a robust functioning regime with both continuous and discrete attractors maintaining grid and assembly scores above 0.8 for at least three consecutive values of *b*_I_. The highest value of *b*_I_ in the functioning regime was labeled as disinhibitory current 0, and we ignored simulations with higher values of *b*_I_. Second, the combination must capture a dysfunctional transition with either the continuous or the discrete attractor dropping its grid or assembly score below 0.2 as *b*_I_ was decreased. The transition was considered achieved once this threshold was crossed, and we ignored simulations with lower values of *b*_I_. Out of the 240 parameter combinations, 58 satisfied both requirements.

### Trained RNNs

#### Network architecture and dynamics

Our RNNs consist of a central recurrent layer of spiking neurons with membrane potentials **v**(*t*) and spike indicators **s**(*t*). Their leaky integrate-and-fire dynamics are implemented based on the snnTorch Python package ^51^. If neuron *i*’s potential exceeds a threshold of 1 at time step *t*, it fires a spike and its potential is reset to 0: *s*_*i*_(*t*) *←* 1 and *v*_*i*_(*t*) *←*0. Otherwise, *s*_*i*_(*t*) = 0.

The recurrent layer neurons receive all-to-all connections from a rate-based input layer **x**(*t*) in addition to their all-to-all recurrent connections:

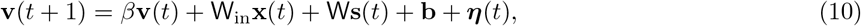

where W_in_ are the input-to-recurrent weights, W are the recurrent-to-recurrent weights, **b** are the recurrent biases, ***η***(*t*) represents independent, zero-mean Gaussian noise, and *β* is the potential decay factor. The recurrent layer neurons send all-to-all connections to a rate-based output layer **y**(*t*):

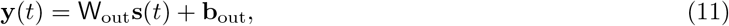

where W_out_ are the recurrent-to-output weights, and **b**_out_ are the output biases.

#### Continuous and discrete tasks

The input and output layers have the same number of neurons *N*_io_ and are organized circularly such that their neurons are located at 1D coordinates *i ∈ {* 1, …, *N*_io_*}* with periodic boundaries. For both continuous and discrete tasks, inputs and targets take the form of a Gaussian bump with fixed width *σ* centered at various positions *z ∈* [0, *N*_io_):

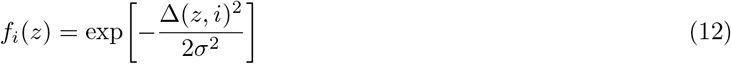

for the periodic 1D distance function

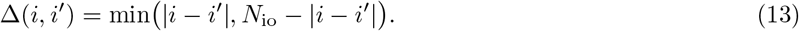

For the continuous task, the target bump **ŷ** is located at the same position as the input bump:

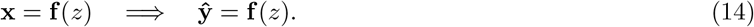

For the discrete task, the target bump is centered at the midpoint of the first half of the output layer if the input bump is centered within the first half of the input layer, and likewise for the second half:

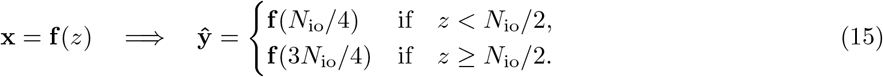

For both tasks, the inputs **x** are presented to the RNN for time steps 1 to 20 and then **x**(*t*) is set to 0. The targets **ŷ** are enforced for time steps 21 to 100 with a mean-squared error loss function:

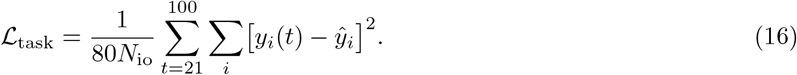

#### Regularizers

To encourage the RNNs to solve continuous and discrete tasks using similar network statistics, we applied regularizers on activities, weights, and biases. We favored a common population activity level *S* in the recurrent layer with

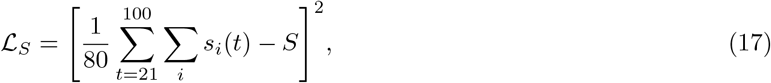

where *s*_*i*_(*t*) is the spike indicator for recurrent neuron *i*. We disfavored concentration of activity in a small number of recurrent neurons with:

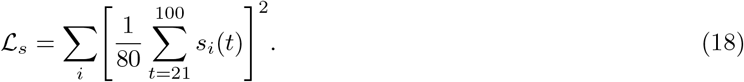

We favored a common mean *μ*_*W*_ and standard deviation *σ*_*W*_ for the recurrent weights with:

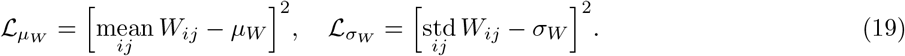

We favored a common mean *μ*_*b*_ and standard deviation *σ*_*b*_ for the recurrent biases with:

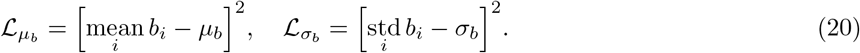

Each of these regularization terms was assigned a strength *λ* to yield the total loss:

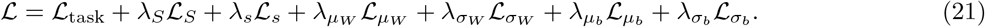

#### Training and evaluation protocols

We trained all weights and biases, which were initialized with PyTorch 2.10.0 default settings. During each training epoch out of 1000 total, we ran the RNN for 100 time steps on a batch of *N*_io_ inputs. We promoted even sampling of input space within each batch by centering one bump at each input neuron and then jittering bump positions by uniformly random values between −0.5 and 0.5. After this forward pass, we computed the total loss Eq. 21 summed over the batch, and we backpropagated errors using the Adam optimizer with learning rate 10^−3^, weight decay 0, *β*_1_ = 0.9, and *β*_2_ = 0.999. To backpropagate through spike discontinuities, we used the fast sigmoid surrogate gradient function with snnTorch 0.9.4 default settings ^51^.

After training, we froze all weights and biases and evaluated these RNNs over 500 time steps with input bumps presented for the first 20 time steps. We used an evaluation batch of *N*_io_ input bumps centered between neurons at positions 0.5, 1.5, …, *N*_io_ − 0.5. Outputs between time steps 21 and 100 are used to compute the test loss, which is identical to the task loss Eq. 16 used during training: *L*_test_ = *L*_task_. Outputs between time steps 101 and 500 are used to compute the generalization loss, which assesses the ability to maintain output bumps at target locations beyond the time horizon experienced during training:

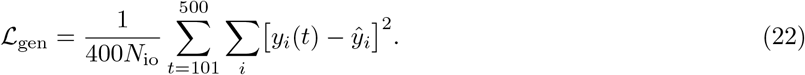

##### Algorithm 1

Spectral circular ordering of recurrent neurons.

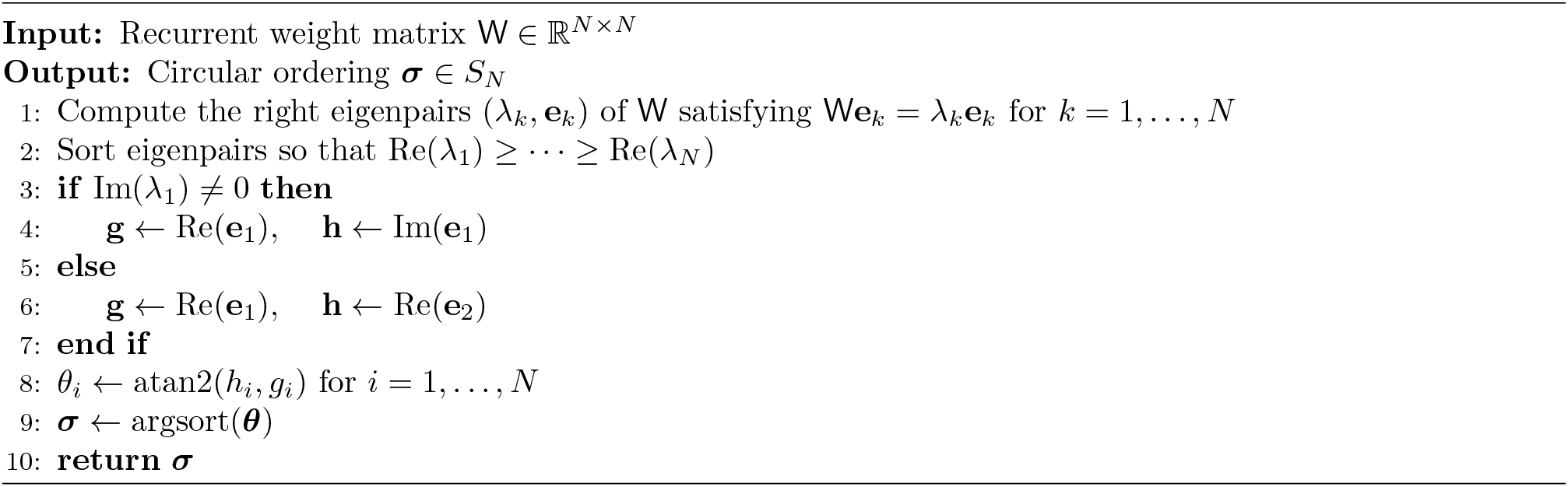

Bias and weight shift perturbations were introduced to the trained RNNs before the start of an evaluation run. We added a small positive constant to either all recurrent biases **b** or all recurrent weights W, and proceeded identically to the unperturbed runs.

Unless otherwise specified, we used standard network hyperparameters provided in Table 2. For the runs in Supp. Fig. 4D, we created discrete tasks with different numbers of targets *N*_target_ by dividing the input and output layer into equally spaced regions and centering target bumps at the midpoint of each region if the input bump is centered within the region:

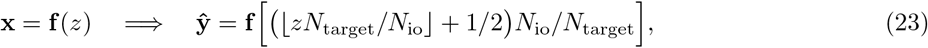

where ⌊·⌋ is the floor function.

**Table 2.** Standard hyperparameters for continuous- and discrete-trained RNNs unless otherwise noted.

| Parameter | Variable | Value |
| --- | --- | --- |
| recurrent layer width | $N$ | 1000 |
| input and output layer width | $N_{\text{io}}$ | 32 |
| potential decay factor | $\beta$ | 0.95 |
| noise magnitude | $\text{std}[\boldsymbol{\eta}]$ | 0.1 |
| input presentation final time step | – | 20 |
| train and test final time step | – | 100 |
| generalization final time step | – | 500 |
| input and target bump width | $\sigma$ | 10 |
| preferred population activity level | $S$ | 12 spk/step |
| preferred weight mean | $\mu_W$ | 0 |
| preferred weight standard deviation | $\sigma_W$ | 0.04 |
| preferred bias mean | $\mu_b$ | 0.01 |
| preferred bias standard deviation | $\sigma_b$ | 0.08 |
| population activity regularizer strength | $\lambda_S$ | $10^{-4}$ |
| single-neuron activity regularizer strength | $\lambda_s$ | 80 |
| weight mean regularizer strength | $\lambda_{\mu_W}$ | 10 |
| weight standard deviation regularizer strength | $\lambda_{\sigma_W}$ | $10^3$ |
| bias mean regularizer strength | $\lambda_{\mu_b}$ | $10^{-2}$ |
| bias standard deviation regularizer strength | $\lambda_{\sigma_b}$ | 1 |
| number of training epochs | – | 1000 |

For the runs in Supp. Fig. 5, we applied more stringent control of network statistics. First, instead of Eq. 17, we used a different population activity regularizer that favors a common population activity level *S* in the recurrent layer at every time step:

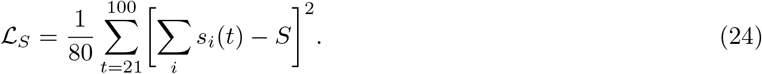

Second, we clamped the diagonal elements of W to 0. These correspond to recurrent self-connections, which effectively set a different reset potential for each neuron, so eliminating them reduces heterogeneity. Third, for evaluation, we used the weights and biases that minimize the loss over the course of training instead of using the final trained network after 1000 epochs. These changes were accompanied by updated regularization hyperparameter values *λ*_*S*_ = 10^−6^, *λ*_*s*_ = 20, *λ*_*μ*_ = 10^3^, *λ*_*σ*_ = 10^4^, *λ*_*μ*_ = 10^2^, *λ*_*σ*_ = 10^3^, *μ*_*b*_ = 0, and *σ*_*b*_ = 0.05.

#### Recurrent layer sorting

We developed a method to order neurons in the recurrent layer based on the learned recurrent weight matrix W after training (Algorithm 1). It is related to spectral clustering ^52^, which uses eigenvalues to group data by similarity, and similar ideas have been used for circular ordering as well as neural connectivity data analysis ^53,54^. It aims to uncover a circular neural ordering in the trained recurrent layer that reflects the circular organization of the input and output layers. We designed the algorithm for the continuous task, since its targets form a circular manifold in state space, but we found that it also recovers meaningful structure in discrete-trained RNNs.

We assume that over the course of training, W has learned to maintain a set of stable states that form a circular manifold in state space. To do so, we expect that it contains the structure of a circulant matrix, like those of continuous ring attractors, but its circular ordering is scrambled by a permutation over neural indices and obscured by noise. We seek to uncover this ordering by inverting the permutation. The eigenvectors of W whose eigenvalues have largest real parts correspond to population activity patterns that dominate the linearized network dynamics. We thus sort the eigenpairs by the real part of their eigenvalues in decreasing order. The eigenvectors of any circulant matrix are the Fourier modes, and a permuted circulant matrix simply has correspondingly permuted Fourier eigenvectors with the same eigenvalues. We expect that the first eigenvector will be the first Fourier mode with a circular topology; the zeroth Fourier mode corresponds to overall changes in activity, and higher order modes correspond to multiple windings around the circle. Thus, we compute the complex argument for each element in the first eigenvector, and their ordering inverts the permutation of the first Fourier mode.

For each Fourier mode in a circulant matrix eigenspectrum except for the zeroth mode and the highest mode of even-sized matrices, there is a corresponding negative mode with the complex conjugate eigenvalue. If the eigenvalue is real, then it is duplicated, and the Fourier modes of this pair are two real sinusoidal functions offset by an arbitrary phase. If this is the case for the first eigenvector, instead of extracting its complex argument, we take the first two eigenvectors and use their element-wise arctangent as the circular coordinate. While we designed this algorithm in the absence of noise, circulant matrices are normal, so their eigenspectra are stable with respect to noise. Thus, our method can recover circular ordering in the presence of small noise. Moreover, due to degeneracies in paired positive and negative Fourier modes, it does not recover the absolute phase offset and orientation of the circular manifold, so we can rotate and reverse the ordering to better align the recurrent layer dynamics with that of the output layer. All recurrent layer spike rasters and weight matrices were sorted using this algorithm before presentation.

### *In vivo* electrophysiology

#### Experimental animals

Adult male wildtype C57BL/6J (C57BL/6JJmsSlc substrain; Japan SLC, Inc.) and CA3-Cre transgenic (C57BL/6-Tg(Grik4-cre)G32-4Stl/J; Jackson Laboratory, RRID: IMSR JAX:006474) mice, 12–24 weeks of age, were used in this study. Mice were bred and genotyped in-house and raised in a temperature- and humidity-controlled room, provided with food and water ad libitum with a 12-h light-dark cycle. All experiments were performed during the light phase. All procedures were approved by the RIKEN Institutional Animal Care and Use Committee and complied with all relevant ethical regulations.

#### Recombinant viruses

The recombinant adeno-associated virus (AAV) vectors used in the experiments were generated in-house. Plasmid pAAV.EF1a.DIO.tetX.2A.mCherry was previously described ^55^. Plasmid pAAV.CaMKII.tetX.2A.mCherry was generated by ligating a PCR insert for the tetX.2A.mCherry open reading frame preceded by a consensus Kozak sequence into the pAAV.CaMKII vector backbone. The pHelper plasmid was purchased from Agilent Technologies (Cat. No. 240071). The pAAV.DJ/8.2.RC plasmid was purchased from Cell Biolabs (Cat. No. VPK-420-DJ-8). All plasmids were sequence-verified.

For AAV production, we used the AAV Helper Free System (Agilent Technologies). The above AAV plasmids were each co-transfected with pHelper (Agilent Technologies) and the pAAV.DJ/8.2.RC (Cell Biolabs) rec/capsid helper plasmid. The trio of plasmids was transfected into the 293FT cell line (Invitrogen) utilizing the 293fectin transfection reagent (Invitrogen). After 72 h, for the AAV.DJ/8.2 serotype AAV vectors, the supernatant was collected and centrifuged at 3000 rpm for 30 min and then filtered through a 0.45-μm filtration unit (Millipore). For the retrograde AAV serotype, the cells and supernatant were collected and put through 3 freeze–thaw cycles (−80°C / 37°) to obtain AAV inside the 293FT cells and then centrifuged at 3000 rpm for 30 min and then filtered through a 0.45-μm filtration unit.

Purification of AAV was carried out by ultracentrifugation (87 000 *g*, 4°C, 2 h) with a 20% sucrose cushion. The supernatant was removed and the pellet was resuspended in phosphate-buffered saline (PBS), aliquoted, and stored at −80°C for long-term storage. The AAV stocks were titered using a custom-ordered AAV stock purchased from Virovek (Hayward, CA) as the reference standard. AAV titer quantification was performed using qPCR with the CFX Opus 96 Real-Time PCR System (Bio-Rad) or StepOne Plus Real Time PCR System (Applied Biosystems), FastStart Universal SYBR Green Master (Roche), and qPCR primers for an approximately 100-bp fragment of the woodchuck hepatitis virus posttranscriptional response element (WPRE) found in all our adeno-associated virus vectors.

#### Virus injection

Mice were deeply anesthetized by a mixture of medetomidine (0.75 mg/kg; Nippon Zenyaku Kogyo), midazolam (4 mg/kg; Sandoz), and butorphanol (5 mg/kg; Meiji Seika Pharma) via i.p. injection and placed in a stereotaxic apparatus (David Kopf Instruments). Anesthetic depth was confirmed by loss of pedal withdrawal reflex, and body temperature was maintained using a heating pad. Following surgery, anesthesia was reversed with i.p. injection of atipamezole hydrochloride (1 mg/kg; Zoetis, Antisedan), and mice were monitored until full recovery. The injection volume and flow rate (50 nL/min) were controlled by an injection pump (Drummond Scientific Company, NANOJECT II) using glass pipettes pulled to an outer tip diameter of 70 μm using a PE-22 puller (Narishige Instruments). Pipettes were removed 10 min after infusion was complete.

pAAV.CaMKII.tetX.2A.mCherry (4.35 *×* 10^12^ viral particles per mL) was injected bilaterally into the MEC (100 nL per injection site, −4.75 mm AP, *±*3.4 mm ML, −2.4 and −3.2 mm DV; −4.6 mm AP, *±*3.4 mm ML, −3.8 and −4.2 mm DV). pAAV.EF1a.DIO.tetX.2A.mCherry (1.57 *×* 10^12^ viral particles per mL) was injected into CA3-Cre mice. Injections were targeted bilaterally to CA3 (300 nL per injection site, −2.04 mm AP, *±*2.35 mm ML, −2.33 mm DV).

#### Probe preparation

Neuropixels 1.0 single-shank and Neuropixels 2.0 four-shank silicon probes (IMEC) were used in this study ^56,57^. Probe tips were sharpened to 20° using a micropipette grinder (Narishige, EG-45) for smooth penetration. Perfluoroalkoxy alkane (PFA)-coated silver wires (AM-Systems, Cat. No. 786500) were soldered to both the reference and ground soldering sites on the flex cable of each probe, while the other side of the silver wire was soldered to a socket pin (Monotaro, PD-1, Cat. No. 00019093). Each soldering point was tested via ohmmeter and then coated with UV-cured resin (BONDIC EVO, BD-SKEJ).

To connect ground pins and reference pins of two probes during the recording, two PFA-coated silver wires (AM-Systems, Cat. No. 786500) were soldered to a gold pin (Inter Medical, IM-C5) as the ground pin, and another two silver wires were soldered to another gold pin as the reference pin. Each soldering point was tested via ohmmeter and then coated with UV-cured resin (BONDIC EVO, BD-SKEJ) to prevent oxidation and reinforce strength. Silicone sealant (World Precision Instruments, KWIK-CAST) was applied at the junction to distribute stress.

Probe insertion trajectories and coordinates were planned, visualized, and optimized using the Neuropixels Trajectory Explorer MATLAB toolbox ^58^ with the Allen Mouse Brain Common Coordinate Framework Atlas ^59^.

#### Probe insertion surgery

Before surgery, the animals received subcutaneous (s.c.) injection of the premedication glycopyrrolate (0.02 mg/kg, s.c.; Selleck, Cat. No. 596-51-0), a quaternary ammonium parasympatholytic, to reduce salivary and bronchial secretions. Then 10 min later anesthesia was induced with urethane (1.3–1.5 g/kg, i.p.; Sigma-Aldrich, Cat. No. 94300). The scalp was shaved and disinfected, and the local anesthetic lidocaine (3 mg/kg, s.c.; Shionogi Pharma) was injected under the scalp. Mice were head-fixed in a dual-arm stereotaxic frame (Kopf Instruments, Model 942), and the eyes were protected with Neo-Medrol EE Ointment (Pfizer). Body temperature was maintained around 37°C using non-electronic, air-activated, disposable heating pads.

After removing the scalp, the skull was disinfected, and tissue adhesive (3M, VetBond) was applied to seal the surrounding tissues and prevent bleeding. Dental cement (Yamahachi Dental) was used to build an approximately 5-mm-high wall around the skull for holding a saline bath during recording. According to planned coordinates, a 2-mm-diameter craniotomy over the right CA3 (center: −2.0 mm AP, 2.0 mm ML), a 1.5-mm-diameter craniotomy over the right MEC (center: −4.70 mm AP, 3.3 mm ML), a 1-mm-diameter craniotomy over the left somatosensory cortex (for reference pin insertion), and a 1-mm-diameter craniotomy over the cerebellum (for ground pin insertion) were carefully performed using stainless punches (Kai Industries, Biopsy Punch). The dura was kept intact to prevent tissue swelling and drift during recording, except for the reference pin, for which the dura was removed to ensure pin insertion into cortex. The craniotomies were covered with gelatin sponge (LTL Pharma, Spongel) to prevent bleeding and maintain moisture for smooth probe penetration.

For probe tracking after the experiment, probes were coated with a solution of DiO (Neuro-DiO, Biotium, Cat. No. 60015) by holding 2 μL in a droplet at the end of a micropipette and touching the droplet along the probe shanks. The probes were mounted to aluminum stereotaxic rods (IMEC) and clamped to motorized micromanipulators (Narishige, MDS-1). For CA3, the Neuropixels 2.0 probe was rotated in azimuth with the inter-shank axis oriented 30° clockwise relative to the ML axis and manually lowered through the center of its craniotomy window until it just penetrated the dura. The micromanipulator was used to insert the probe at 2 μm/s to a depth of 3000 μm. For MEC, the Neuropixels 1.0 probe was rotated in elevation with the tip pointing 6° anteriorly relative to the DV axis and manually lowered through its craniotomy window directly anterior to the transverse sinus until it just penetrated the dura. The micromanipulator was used to insert the probe at 2 μm/s until the shank was observed to bend slightly as an indication of hitting the skull beneath, after which the probe was retracted by about 20 μm until it became straight again ^60^.

The reference wires of each probe were connected to one gold pin and inserted approximately 1–2 mm inside the somatosensory cortex through the reference craniotomy, and the ground wires of each probe were connected to another gold pin and inserted above the cerebellum through the ground craniotomy. Saline was added inside the dental cement wall to immerse the probe shanks, reference pin, and ground pin.

#### Recording protocol

Probes were configured, synchronized, visualized, and streamed to disk using a PXIe-PCIe interface (National Instruments, PCIe-8382) and chassis (National Instruments, PXIe-1071) through SpikeGLX software ^61^. For MEC, we used the default single-shank Neuropixels 1.0 channel map, which selects its bottom channel bank. For CA3, we selected multi-shank Neuropixels 2.0 channels estimated to best capture the CA3 pyramidal layer based on electrophysiological signatures.

Recording consisted of a 40-min baseline period followed by a low-dose PTZ injection (30 mg/kg, i.p.; Selleck, Cat. No. 54-95-50) and 50 min of subsequent recording. Acquisition was then paused, and the AP filter was turned off and the gain was lowered to 125 for the Neuropixels 1.0 probe to acquire broadband signal at high sample rate similarly to the Neuropixels 2.0 probe. High-dose PTZ was injected (150 mg/kg, i.p.) followed by 60 min of subsequent recording.

#### Histology and immunohistochemistry

After recording completion, the animals were deeply anesthetized with urethane (Sigma-Aldrich, Cat. No. 94300) and transcardially perfused with 0.1 M phosphate-buffered saline (PBS, pH 7.4; Nacalai Tesque, Cat. No. 05150-45), followed by 4% paraformaldehyde (PFA; Nacalai Tesque, Cat. No. 09154-85) in PBS. The brains were removed and postfixed overnight at 4°C with 4% PFA. Sagittal sections of 60 μm thickness were prepared using a vibratome (Leica), collected in 24-well plates with antibacterial solution (PBS + 0.05% sodium azide; Nacalai Tesque, Cat. No. 26628-22-8), and stored at 4°C.

For viral expression verification and probe tracking, slices were washed in PBS and then mounted with VECTASHIELD Mounting Medium with DAPI (Vector Laboratories, Cat. No. H-1200, RRID: AB 2336790). Probe tracks marked by DiO were anatomically aligned with the AP histology MATLAB toolbox ^62^ to obtain region estimates for each probe channel. These labels were reconciled with low-frequency (1–500 Hz) and high-frequency (0.3–30 kHz) root-mean-square LFP amplitudes and spike locations estimated by Kilosort (described below). This combination of histological channel labels and electrophysiological channel properties was used to select channels located in MEC and CA3 (Supp. Fig. 6).

For VAMP2 (synaptobrevin-2) staining to confirm region-specific cleavage by expressed tetanus toxin, sections were first blocked in PBS + 0.03% Triton X-100 + 3% normal donkey serum, incubated overnight in a primary antibody prepared in blocking solution (rabbit anti-VAMP2, 1:250; Synaptic Systems, Cat. No. 104 202), followed by three washes with PBS + 0.03% Triton X-100 (10 min each) and subsequently incubated with Alexa Fluor 488-conjugated donkey anti-rabbit secondary antibody (1:500; Thermo Fisher Scientific, Cat. No. A32790TR, RRID: AB 2866495) prepared in blocking solution for 2 h at room temperature. Following three washes with PBS + 0.03% Triton X-100 (10 min each), sections were mounted with VECTASHIELD Mounting Medium with DAPI (Vector Laboratories, Cat. No. H-1200, RRID: AB 2336790). The VAMP2 optical signal was quantified for each mouse by first identifying the following regions in four sections: CA1 striatum radiatum, CA1 stratum lacunosum-moleculare, dentate gyrus (DG) outer molecular layer, DG middle molecular layer, and DG inner molecular layer. Fluorescence pixel values for each section were divided by the mean value of the subiculum, and these normalized pixel values were averaged within and across sections for each region (Supp. Fig. 7).

#### Spike sorting

Extracellular spike waveforms were detected and clustered into putative single-neuron spike trains using the Kilosort 4.1.1 and SpikeInterface 0.103.2 Python packages ^63,64^. Neuropixels recordings from recording start to 50 min after low-dose PTZ injection were extracted and then preprocessed with phase correction, global common average referencing, and highpass filtering above 300 Hz. Kilosort4 was run with max peels = 200, which increases the number of spike detection iterations as previously suggested for MEC recordings ^19^.

The inter-spike interval (ISI) violation ratio computed with the SpikeInterface compute isi violations function was primarily used to assess the quality of each cluster. It was computed using a permissive ISI threshold of 1 ms to allow for more rapid spiking dynamics under PTZ perturbations. Clusters with ISI violation ratio below 0.2 were considered to represent single units and accepted for further analysis. All other clusters were subjected to an incremental pruning of spikes with outlying waveforms as follows. For each cluster, the spatiotemporal profile of each spike was projected into a 4-dimensional space consisting of the top 2 waveform PCs of the 2 channels with largest trough-to-peak template amplitudes. Each dimension was *z*-scored, all points with distance greater than 8 from the origin were excluded, and remaining points were whitened. These manipulations represent a linear transformation of spike waveform shape into a standardized space. Each cluster was then incrementally pruned by removing, in increasing percentages from 1% to 30%, the spikes furthest from the origin in this standardized space. At each step, the ISI violation ratio of the remaining spikes was recomputed, and if it decreased below 0.2, then the pruning process was terminated and the resulting cluster was accepted for further analysis. Clusters whose ISI violation ratio never fell below 0.2 under pruning were excluded.

All accepted clusters were manually curated with the phy 2.0 Python package ^65^ to exclude clusters with markedly artifactual waveform shapes and to split and merge clusters where clearly appropriate. All clusters with firing rates below 0.2 Hz were excluded.

#### LFP analysis

To obtain the results in Fig. 4, the LFP after high-dose PTZ injection was broadband-filtered between 1–500 Hz and downsampled to 1000 Hz. For visualization, 5–20 evenly spaced channels spanning the recorded areas of MEC and CA3 were selected. For LFP peak amplitude analysis, 40 evenly spaced channels spanning the recorded areas of MEC and CA3 were selected. The Hilbert transform was applied to the signal in each channel between 30–60 min after high-dose PTZ injection, and its complex magnitude was computed to yield the Hilbert amplitude. The 95th percentile of the Hilbert amplitude over time was defined to be the peak amplitude of each LFP channel. For band power analysis in Supp. Fig. 9, 40 evenly spaced channels spanning the recorded areas of MEC and CA3 were selected. The Welch periodogram method for spectral density estimation was applied to each channel over each 2-min non-overlapping window between 30–60 min after high-dose PTZ injection. A Hann window function was used with segment length 2 s and overlap length 1 s. Power spectral densities within each frequency band were averaged to yield band power.

We computed the Granger directionality index using the LFP signals of all MEC and CA3 channels. We extracted the signal between 30–60 min after high-dose PTZ injection using 1-second timestamps provided by the sync signal common to both Neuropixels probes. This allows for sub-millisecond temporal alignment between MEC and CA3 signals, which is necessary for analyzing their temporal relationship at the millisecond level. They were broadband-filtered between 1–500 Hz, downsampled to 1000 Hz, and processed to yield Hilbert amplitudes. For computational tractability, these Hilbert amplitude signals were downsampled to 500 Hz and projected, separately for each brain region, along their leading principal components (PCs) that explained 99% cumulative variance. Following methods implemented in the statsmodels Python package, these PCs were fit to linear vector autoregressive models with 10 time lags of 2 ms. For each brain region, a restricted and an unrestricted model were fit, and their sum of squared residuals were used to calculate Granger causality *F*-statistics for MEC Granger-causing CA3 (*F*_MEC*→*CA3_) and CA3 Granger-causing MEC (*F*_CA3*→*MEC_). Because the degrees of freedom for both causal directions are equal, these *F*-statistics could then be combined into a directionality index ^66,67^

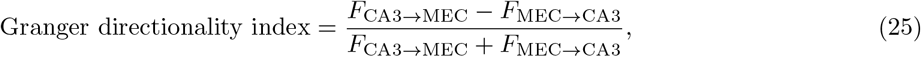

which ranges from −1 to 1. Negative values indicate that predominant temporal ordering is MEC leading CA3, and positive values indicate the reverse. In Fig. 4J,K, separate models were fit for each 1-min non-overlapping window between 30–60 min, and in Fig. 4L, one set of models was fit for the entire interval 30–60 min.

#### Single-unit analysis

For the results in Fig. 5, we identified neurons to be the putative single-unit spike clusters accepted after spike sorting, automated quality assessment with spike pruning, manual curation, and selection of channels within MEC and CA3. In Figs. 5E and S10, neurons in each region were sorted using the Rastermap 1.0 Python package ^68^. In Fig. 5F, normalized population activity was defined to be the mean firing rate in each region between 10–40 min after low-dose PTZ injection divided by its value between 0–30 min of baseline recording.

We computed temporal continuity by partitioning the interval between 10–40 min after low-dose PTZ injection into 5-min subepochs to accommodate units that emerge into or drop out of the recording. For each subepoch and region, neurons with time-averaged firing rate below 0.05 Hz or above the 90th percentile were excluded. To prevent differences in population size from confounding our results, 20 subsampled sets of 30 randomly selected neurons were formed. In each subsample, firing rates within 200-ms bins were computed to yield the *network state*, a temporally evolving 30-dimensional vector of neural activity. Time bins with population-averaged firing rate below 0.01 Hz or above the 90th percentile were excluded. To compensate for any slow changes in overall neural activity due to PTZ absorption, the firing rate of each neuron was detrended by subtracting its Gaussian-filtered value using a standard deviation of 30 s, which yields the detrended network state **∑** (*t*). The temporal continuity for time bin *t* is then

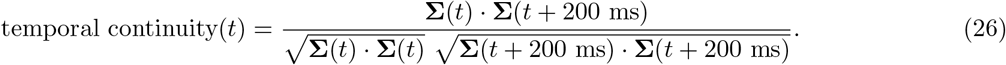

In Fig. 5G–I, the temporal continuity was computed for 10 randomly selected time bins per subepoch and subsample, and the results were aggregated across subepochs and subsamples. For detrended cosine similarities at other time lags in Supp. Fig. 11, the values of 200 ms in Eq. 26 were replaced by the corresponding lag value. For the attractor simulation results in Fig. 5C,D, temporal continuity was computed similarly to the case of experimental recordings except that the entire 2-min simulation after a 3-s transient was used without forming subepochs.

## Acknowledgments

The authors acknowledge Yinghao Liu, Eric T. N. Overton, and Yusuke Kasuga for their assistance and advice on experimental protocols and analysis. They also acknowledge Naoyo Horiguchi and Michiko Fujisawa for administrative and logistical support.

## Funding

Japan Society for the Promotion of Science 22K15209: LK

Ministry of Education, Culture, Sports, Science and Technology (Japan) 23H05478: TJM RIKEN Center for Brain Science: TJM, LK

## Author contributions

ML, SE, and IR were respectively the primary contributors to the animal experiments, attractor networks, and trained RNNs involved in this work.

Conceptualization: LK

Methodology: ML, SE, IR, DP, TJM, LK Investigation – experimental: ML, LK Investigation – computational: SE, IR, IT, HN, LK Resources: DP, AJYH, TJM, LK

Visualization: ML, SE, IR, LK

Writing – original draft: ML, SE, IR, LK

Writing – review & editing: ML, SE, IR, DP, AJYH, IT, HN, DY, TJM, LK Supervision: DY, TJM, LK

## Data and code availability

Data and code will be made publicly available upon acceptance for publication.

## References

[1] O. Devinsky, A. Vezzani, T. J. O’Brien, N. Jette, I. E. Scheffer, M. de Curtis, and P. Perucca. Epilepsy. Nat. Rev. Dis. Primers, 4:18024, 2018. doi: 10.1038/nrdp.2018.24.

[2] C. Waruiru and R. Appleton. Febrile seizures: an update. Arch. Dis. Child., 89:751–756, 2004. doi: 10.1136/adc.2003.028449.

[3] H.-Y. Chen, T. E. Albertson, and K. R. Olson. Treatment of drug-induced seizures: Treatment of drug-induced seizures. Br. J. Clin. Pharmacol., 81:412–419, 2016. doi: 10.1111/bcp.12720.

[4] R. Nardone, F. Brigo, and E. Trinka. Acute symptomatic seizures caused by electrolyte disturbances. J. Clin. Neurol., 12:21–33, 2016. doi: 10.3988/jcn.2016.12.1.21.

[5] A. J. Trevelyan, D. Sussillo, B. O. Watson, and R. Yuste. Modular propagation of epileptiform activity: evidence for an inhibitory veto in neocortex. J. Neurosci., 26:12447–12455, 2006. doi: 10.1523/JNEUROSCI.2787-06.2006.

[6] K. P. Lillis, M. A. Kramer, J. Mertz, K. J. Staley, and J. A. White. Pyramidal cells accumulate chloride at seizure onset. Neurobiol. Dis., 47:358–366, 2012. doi: 10.1016/j.nbd.2012.05.016.

[7] V. K. Jirsa, W. C. Stacey, P. P. Quilichini, A. I. Ivanov, and C. Bernard. On the nature of seizure dynamics. Brain, 137:2210–2230, 2014. doi: 10.1093/brain/awu133.

[8] L. M. Y. Yu, D. Polygalov, M. E. Wintzer, M.-C. Chiang, and T. J. McHugh. CA3 synaptic silencing attenuates kainic acid-induced seizures and hippocampal network oscillations. eNeuro, 3, 2016. doi: 10.1523/ENEURO.0003-16.2016.

[9] M. Wenzel, J. P. Hamm, D. S. Peterka, and R. Yuste. Acute focal seizures start as local synchronizations of neuronal ensembles. J. Neurosci., 39:8562–8575, 2019. doi: 10.1523/JNEUROSCI.3176-18.2019.

[10] J.-Y. Liou, E. H. Smith, L. M. Bateman, S. L. Bruce, G. M. McKhann, R. R. Goodman, R. G. Emerson, C. A. Schevon, and L. F. Abbott. A model for focal seizure onset, propagation, evolution, and progression. Elife, 9:e50927, 2020. doi: 10.7554/eLife.50927.

[11] D. Hadjiabadi, M. Lovett-Barron, I. G. Raikov, F. T. Sparks, Z. Liao, S. C. Baraban, J. Leskovec, A. Losonczy, K. Deisseroth, and I. Soltesz. Maximally selective single-cell target for circuit control in epilepsy models. Neuron, 109:2556–2572.e6, 2021. doi: 10.1016/j.neuron.2021.06.007.

[12] G. D. Rosen, A. G. Williams, J. A. Capra, M. T. Connolly, B. Cruz, L. Lu, D. C. Airey, K. Kulkarni, and R. W. Williams. The mouse brain library @ www.mbl.org. Int Mouse Genome Conference, 14:166, 2000.

[13] D. A. McCormick and D. Contreras. On the cellular and network bases of epileptic seizures. Annu. Rev. Physiol., 63:815–846, 2001. doi: 10.1146/annurev.physiol.63.1.815.

[14] W. Tatum. Mesial temporal lobe epilepsy. Journal of Clinical Neurophysiology, 29:356–365, 2012. doi: 10.1097/wnp.0b013e31826b3ab7.

[15] D. Marr. Simple memory: a theory for archicortex. Philos. Trans. R. Soc. Lond. B Biol. Sci., 262:23–81, 1971. doi: 10.1098/rstb.1971.0078.

[16] I. Zutshi, M. L. Fu, V. Lilascharoen, J. K. Leutgeb, B. K. Lim, and S. Leutgeb. Recurrent circuits within medial entorhinal cortex superficial layers support grid cell firing. Nat. Commun., 9:3701, 2018. doi: 10.1038/s41467-018-06104-5.

[17] K. Yoon, M. A. Buice, C. Barry, R. Hayman, N. Burgess, and I. R. Fiete. Specific evidence of low-dimensional continuous attractor dynamics in grid cells. Nat. Neurosci., 16:1077–1084, 2013. doi: 10.1038/nn.3450.

[18] Y. Gu, S. Lewallen, A. A. Kinkhabwala, C. Domnisoru, K. Yoon, J. L. Gauthier, I. R. Fiete, and D. W. Tank. A map-like micro-organization of grid cells in the medial entorhinal cortex. Cell, 175:736–750.e30, 2018. doi: 10.1016/j.cell.2018.08.066.

[19] R. J. Gardner, E. Hermansen, M. Pachitariu, Y. Burak, N. A. Baas, B. A. Dunn, M.-B. Moser, and E. I. Moser. Toroidal topology of population activity in grid cells. Nature, 602:123–128, 2022. doi: 10.1038/s41586-021-04268-7.

[20] A. Treves and E. T. Rolls. Computational analysis of the role of the hippocampus in memory. Hippocampus, 4:374–391, 1994. doi: 10.1002/hipo.450040319.

[21] T. Sasaki, N. Matsuki, and Y. Ikegaya. Metastability of active CA3 networks. Journal of Neuroscience, 27:517–528, 2007. doi: 10.1523/jneurosci.4514-06.2007.

[22] T. D. Goode, K. Z. Tanaka, A. Sahay, and T. J. McHugh. An integrated index: Engrams, place cells, and hippocampal memory. Neuron, 107:805–820, 2020. doi: 10.1016/j.neuron.2020.07.011.

[23] R. P. Sammons, M. Vezir, L. Moreno-Velasquez, G. Cano, M. Orlando, M. Sievers, E. Grasso, V. D. Metodieva, R. Kempter, H. Schmidt, and D. Schmitz. Structure and function of the hippocampal CA3 module. Proc. Natl. Acad. Sci. U. S. A., 121:e2312281120, 2024. doi: 10.1073/pnas.2312281120.

[24] Y. Li, J. J. Briguglio, S. Romani, and J. C. Magee. Mechanisms of memory-supporting neuronal dynamics in hippocampal area CA3. Cell, 188:868, 2025. doi: 10.1016/j.cell.2025.01.020.

[25] J. Widloski and I. R. Fiete. A model of grid cell development through spatial exploration and spike time-dependent plasticity. Neuron, 83:481–495, 2014. doi: 10.1016/j.neuron.2014.06.018.

[26] L. Kang and M. R. DeWeese. Replay as wavefronts and theta sequences as bump oscillations in a grid cell attractor network. Elife, 8:e46351, 2019. doi: 10.7554/elife.46351.

[27] R. L. Macdonald and J. L. Barker. Pentylenetetrazol and penicillin are selective antagonists of GABA-mediated post-synaptic inhibition in cultured mammalian neurones. Nature, 267:720–721, 1977. doi: 10.1038/267720a0.

[28] C. J. Yuskaitis, L.-A. Rossitto, K. J. Groff, S. C. Dhamne, B. Zhang, L. K. Lalani, A. K. Singh, A. Rotenberg, and M. Sahin. Factors influencing the acute pentylenetetrazole-induced seizure paradigm and a literature review. Ann. Clin. Transl. Neurol., 8:1388–1397, 2021. doi: 10.1002/acn3.51375.

[29] K. F. Green and J. N. Rawlins. Hippocampal theta in rats under urethane: generators and phase relations. Electroencephalogr. Clin. Neurophysiol., 47:420–429, 1979. doi: 10.1016/0013-4694(79)90158-5.

[30] A. Czurkó, J. Huxter, Y. Li, B. Hangya, and R. U. Muller. Theta phase classification of interneurons in the hippocampal formation of freely moving rats. J. Neurosci., 31:2938–2947, 2011. doi: 10.1523/JNEUROSCI.5037-10.2011.

[31] G. Buzsáki. Hippocampal sharp wave-ripple: A cognitive biomarker for episodic memory and planning. Hippocampus, 25:1073–1188, 2015. doi: 10.1002/hipo.22488.

[32] J. P. Dreier and U. Heinemann. Regional and time dependent variations of low Mg2+ induced epileptiform activity in rat temporal cortex slices. Exp. Brain Res., 87:581–596, 1991. doi: 10.1007/bf00227083.

[33] M. Barbarosie and M. Avoli. CA3-driven hippocampal-entorhinal loop controls rather than sustains in vitro limbic seizures. Journal of Neuroscience, 17:9308–9314, 1997. doi: 10.1523/jneurosci.17-23-09308.1997.

[34] S. S. Spencer and D. D. Spencer. Entorhinal-hippocampal interactions in medial temporal lobe epilepsy. Epilepsia, 35:721–727, 1994. doi: 10.1111/j.1528-1157.1994.tb02502.x.

[35] B. A. Assaf and J. S. Ebersole. Continuous source imaging of scalp ictal rhythms in temporal lobe epilepsy. Epilepsia, 38:1114–1123, 1997. doi: 10.1111/j.1528-1157.1997.tb01201.x.

[36] N. Bernasconi, A. Bernasconi, F. Andermann, F. Dubeau, W. Feindel, and D. C. Reutens. Entorhinal cortex in temporal lobe epilepsy: a quantitative MRI study: A quantitative MRI study. Neurology, 52: 1870–1876, 1999. doi: 10.1212/wnl.52.9.1870.

[37] P. A. Schwartzkroin and D. A. Prince. Cellular and field potential properties of epileptogenic hippocampal slices. Brain Res., 147:117–130, 1978. doi: 10.1016/0006-8993(78)90776-x.

[38] A. Agmon and J. E. Wells. The role of the hyperpolarization-activated cationic current I(h) in the timing of interictal bursts in the neonatal hippocampus. J. Neurosci., 23:3658–3668, 2003. doi: 10.1523/jneurosci.23-09-03658.2003.

[39] R. D. Traub and R. K. Wong. Cellular mechanism of neuronal synchronization in epilepsy. Science, 216: 745–747, 1982. doi: 10.1126/science.7079735.

[40] T. I. Netoff, R. Clewley, S. Arno, T. Keck, and J. A. White. Epilepsy in small-world networks. J. Neurosci., 24:8075–8083, 2004. doi: 10.1523/JNEUROSCI.1509-04.2004.

[41] C. A. Schevon, S. Tobochnik, T. Eissa, E. Merricks, B. Gill, R. R. Parrish, L. M. Bateman, G. M. McKhann, Jr, R. G. Emerson, and A. J. Trevelyan. Multiscale recordings reveal the dynamic spatial structure of human seizures. Neurobiol. Dis., 127:303–311, 2019. doi: 10.1016/j.nbd.2019.03.015.

[42] S. A. Moosavi, V. K. Jirsa, and W. Truccolo. Critical dynamics in the spread of focal epileptic seizures: Network connectivity, neural excitability and phase transitions. PLoS One, 17:e0272902, 2022. doi: 10.1371/journal.pone.0272902.

[43] Y. Feng, K. S. Diego, Z. Dong, Z. Christenson Wick, L. Page-Harley, V. Page-Harley, J. Schnipper, S. I. Lamsifer, Z. T. Pennington, L. M. Vetere, P. A. Philipsberg, I. Soler, A. Jurkowski, C. J. Rosado, N. N. Khan, D. J. Cai, and T. Shuman. Distinct changes to hippocampal and medial entorhinal circuits emerge across the progression of cognitive deficits in epilepsy. Cell Rep., 44:115131, 2025. doi: 10.1016/j.celrep.2024.115131.

[44] P. R. Montague, R. J. Dolan, K. J. Friston, and P. Dayan. Computational psychiatry. Trends Cogn. Sci., 16:72–80, 2012. doi: 10.1016/j.tics.2011.11.018.

[45] X.-J. Wang and J. H. Krystal. Computational psychiatry. Neuron, 84:638–654, 2014. doi: 10.1016/j.neuron.2014.10.018.

[46] Q. J. M. Huys, T. V. Maia, and M. J. Frank. Computational psychiatry as a bridge from neuroscience to clinical applications. Nat. Neurosci., 19:404–413, 2016. doi: 10.1038/nn.4238.

[47] G. J. Balas and J. C. Doyle. Robustness and performance trade-offs in control design for flexible structures. IEEE Trans. Control Syst. Technol., 2:352–361, 1994. doi: 10.1109/87.338656.

[48] R. Kubo. The fluctuation-dissipation theorem. Rep. Prog. Phys., 29:255–284, 1966. doi: 10.1088/0034-4885/29/1/306.

[49] G. Wang. A fast MCMC algorithm for the uniform sampling of binary matrices with fixed margins. Electron. J. Stat., 14:1690–1706, 2020. doi: 10.1214/20-ejs1702.

[50] L. Kang and V. Balasubramanian. A geometric attractor mechanism for self-organization of entorhinal grid modules. Elife, 8:e46687, 2019. doi: 10.7554/elife.46687.

[51] J. K. Eshraghian, M. Ward, E. O. Neftci, X. Wang, G. Lenz, G. Dwivedi, M. Bennamoun, D. S. Jeong, and W. D. Lu. Training spiking neural networks using lessons from deep learning. Proc. IEEE Inst. Electr. Electron. Eng., 111:1016–1054, 2023. doi: 10.1109/jproc.2023.3308088.

[52] U. von Luxburg. A tutorial on spectral clustering. Stat. Comput., 17:395–416, 2007. doi: 10.1007/s11222-007-9033-z.

[53] A. Recanati, T. Kerdreux, and A. d’Aspremont. Reconstructing latent orderings by spectral clustering. ArXiv, 2018. doi: 10.48550/arXiv.1807.07122.

[54] D. A. Pospisil, M. J. Aragon, S. Dorkenwald, A. Matsliah, A. R. Sterling, P. Schlegel, S.-C. Yu, C. E. McKellar, M. Costa, K. Eichler, G. S. X. E. Jefferis, M. Murthy, and J. W. Pillow. The fly connectome reveals a path to the effectome. Nature, 634:201–209, 2024. doi: 10.1038/s41586-024-07982-0.

[55] R. Boehringer, D. Polygalov, A. J. Y. Huang, S. J. Middleton, V. Robert, M. E. Wintzer, R. A. Piskorowski, V. Chevaleyre, and T. J. McHugh. Chronic loss of CA2 transmission leads to hippocampal hyperexcitability. Neuron, 94:642–655.e9, 2017. doi: 10.1016/j.neuron.2017.04.014.

[56] J. J. Jun, N. A. Steinmetz, J. H. Siegle, D. J. Denman, M. Bauza, B. Barbarits, A. K. Lee, C. A. Anastassiou, A. Andrei, c. Aydin, M. Barbic, T. J. Blanche, V. Bonin, J. a. Couto, B. Dutta, S. L. Gratiy, D. A. Gutnisky, M. Häusser, B. Karsh, P. Ledochowitsch, C. M. Lopez, C. Mitelut, S. Musa, M. Okun, M. Pachitariu, J. Putzeys, P. D. Rich, C. Rossant, W.-L. Sun, K. Svoboda, M. Carandini, K. D. Harris, C. Koch, J. O’Keefe, and T. D. Harris. Fully integrated silicon probes for high-density recording of neural activity. Nature, 551:232–236, 2017. doi: 10.1038/nature24636.

[57] N. A. Steinmetz, C. Aydin, A. Lebedeva, M. Okun, M. Pachitariu, M. Bauza, M. Beau, J. Bhagat, C. Böhm, M. Broux, S. Chen, J. Colonell, R. J. Gardner, B. Karsh, F. Kloosterman, D. Kostadinov, C. Mora-Lopez, J. O’Callaghan, J. Park, J. Putzeys, B. Sauerbrei, R. J. J. van Daal, A. Z. Vollan, S. Wang, M. Welkenhuysen, Z. Ye, J. T. Dudman, B. Dutta, A. W. Hantman, K. D. Harris, A. K. Lee, E. I. Moser, J. O’Keefe, A. Renart, K. Svoboda, M. Häusser, S. Haesler, M. Carandini, and T. D. Harris. Neuropixels 2.0: A miniaturized high-density probe for stable, long-term brain recordings. Science, 372:eabf4588, 2021. doi: 10.1126/science.abf4588.

[58] A. Peters. petersaj/neuropixels trajectory explorer: Alternate probe support, 2024. doi: 10.5281/ZENODO.10793946.

[59] Q. Wang, S.-L. Ding, Y. Li, J. Royall, D. Feng, P. Lesnar, N. Graddis, M. Naeemi, B. Facer, A. Ho, T. Dolbeare, B. Blanchard, N. Dee, W. Wakeman, K. E. Hirokawa, A. Szafer, S. M. Sunkin, S. W. Oh, A. Bernard, J. W. Phillips, M. Hawrylycz, C. Koch, H. Zeng, J. A. Harris, and L. Ng. The allen mouse brain common coordinate framework: A 3D reference atlas. Cell, 181:936–953.e20, 2020. doi: 10.1016/j.cell.2020.04.007.

[60] C. S. Mallory, K. Hardcastle, M. G. Campbell, A. Attinger, I. I. C. Low, J. L. Raymond, and L. M. Giocomo. Mouse entorhinal cortex encodes a diverse repertoire of self-motion signals. Nat. Commun., 12: 671, 2021. doi: 10.1038/s41467-021-20936-8.

[61] J. Putzeys, B. C. Raducanu, A. Carton, J. De Ceulaer, B. Karsh, J. H. Siegle, N. Van Helleputte, T. D. Harris, B. Dutta, S. Musa, and C. Mora Lopez. Neuropixels data-acquisition system: A scalable platform for parallel recording of 10 000+ electrophysiological signals. IEEE Trans. Biomed. Circuits Syst., 13: 1635–1644, 2019. doi: 10.1109/TBCAS.2019.2943077.

[62] A. Peters. Petersaj/AP histology: AP histology v2.0.0, 2026. doi: 10.5281/ZENODO.18746423.

[63] M. Pachitariu, S. Sridhar, J. Pennington, and C. Stringer. Spike sorting with Kilosort4. Nat. Methods, 21: 914–921, 2024. doi: 10.1038/s41592-024-02232-7.

[64] A. P. Buccino, C. L. Hurwitz, S. Garcia, J. Magland, J. H. Siegle, R. Hurwitz, and M. H. Hennig. SpikeInterface, a unified framework for spike sorting. Elife, 9:e61834, 2020. doi: 10.7554/eLife.61834.

[65] C. Rossant and K. D. Harris. Hardware-accelerated interactive data visualization for neuroscience in python. Front. Neuroinform., 7:36, 2013. doi: 10.3389/fninf.2013.00036.

[66] A. Roebroeck, E. Formisano, and R. Goebel. Mapping directed influence over the brain using granger causality and fMRI. Neuroimage, 25:230–242, 2005. doi: 10.1016/j.neuroimage.2004.11.017.

[67] M. Vinck, L. Huurdeman, C. A. Bosman, P. Fries, F. P. Battaglia, C. M. A. Pennartz, and P. H. Tiesinga. How to detect the granger-causal flow direction in the presence of additive noise? Neuroimage, 108:301–318, 2015. doi: 10.1016/j.neuroimage.2014.12.017.

[68] C. Stringer, L. Zhong, A. Syeda, F. Du, M. Kesa, and M. Pachitariu. Rastermap: a discovery method for neural population recordings. Nat. Neurosci., 28:201–212, 2025. doi: 10.1038/s41593-024-01783-4.

