## Supplementary Figures for "Computational demands shape seizure susceptibility in recurrent neural networks"

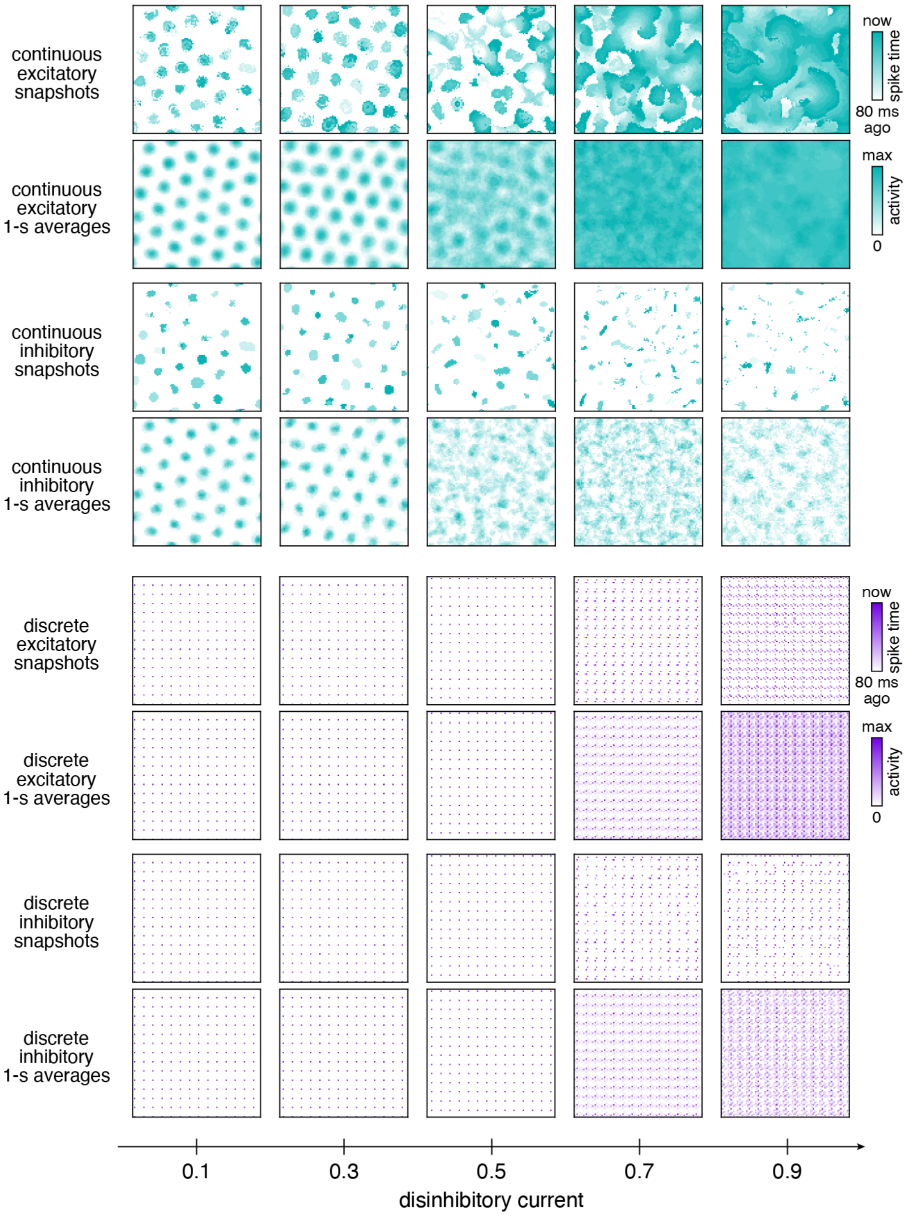


**Supplementary Figure 1**: Extended results for the continuous and discrete attractor networks in Fig. 2 of the main text. Excitatory and inhibitory neuron activity over a range of disinhibitory currents. Each pixel corresponds to one neuron.

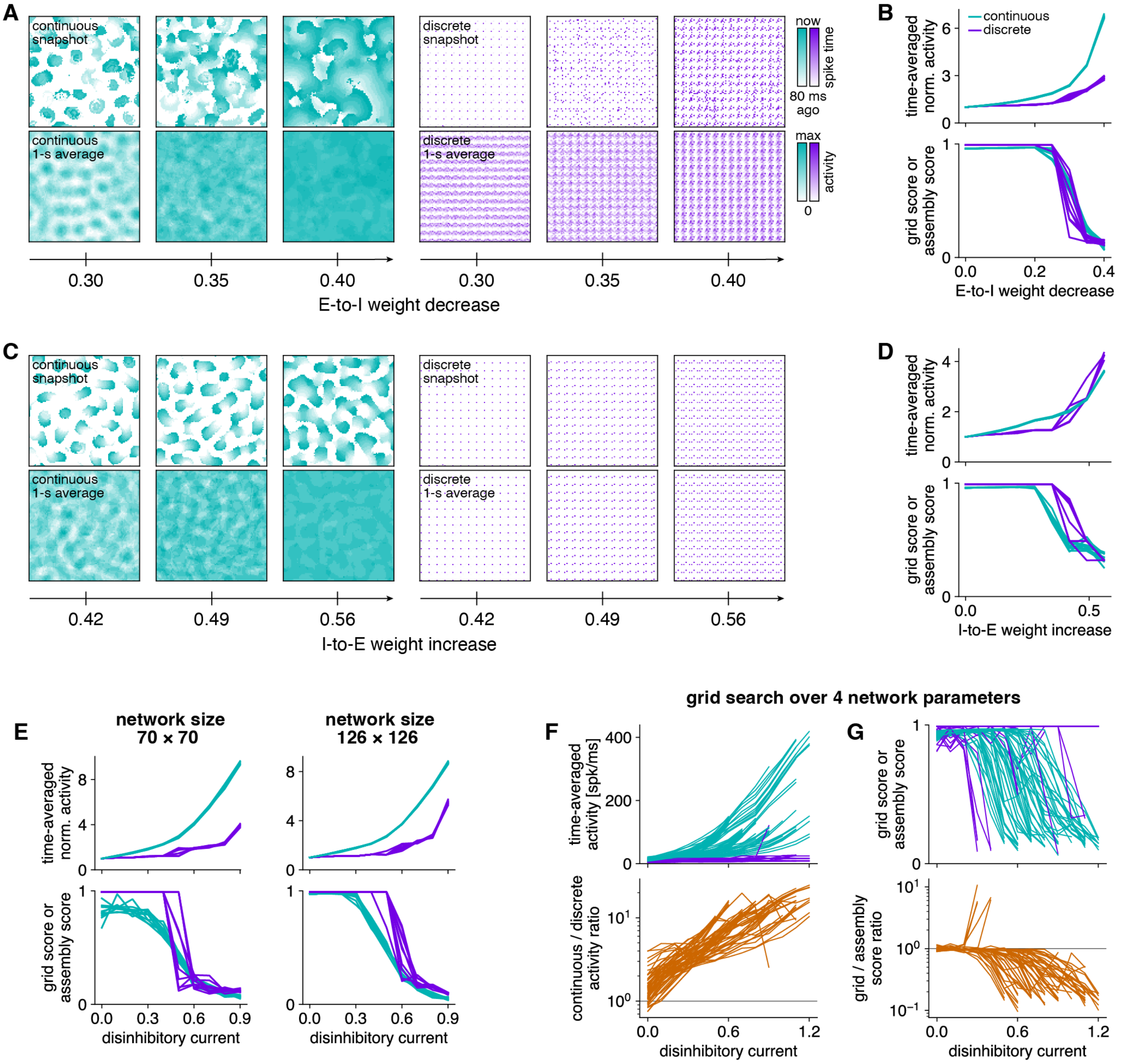


**Supplementary Figure 2**: Continuous and discrete attractor networks with different perturbations and parameters. (**A**,**B**) Perturbing networks by decreasing the synaptic weights from excitatory (E) to inhibitory (I) neurons. (**A**) Excitatory neuron activity over a range of E-to-I weight decreases. Each pixel corresponds to one neuron. (**B**) Top, time-averaged population activity divided by its unperturbed value, and bottom, grid score for continuous attractors and assembly score for discrete attractors. Data from 10 replicate simulations, and each line corresponds to one replica. (**C**,**D**) Similar to **A**,**B**, but for increasing I-to-E synaptic weights. (**E**) Similar to **B**, but for smaller and larger network sizes compared to 98 $\times$ 98 used in Fig. 2 of the main text. (**F**,**G**) Results for paired continuous and discrete attractor networks perturbed with disinhibitory current over parameters found by a grid search over excitatory drive, E-to-E synaptic weight, E-to-I synaptic weight, and I-to-E synaptic weight. The search found 58 parameter combinations with functioning continuous and discrete attractors at low disinhibitory current and dysfunctional continuous or discrete attractors at high disinhibitory current. (**F**) Top, time-averaged population activity, and bottom, ratio between paired continuous and discrete attractors with the same parameters. Each line corresponds to one parameter combination. (**G**) Top, grid score for continuous attractors and assembly score for discrete attractors, and bottom, ratio between paired continuous and discrete networks with the same parameters. Each line corresponds to one parameter combination.

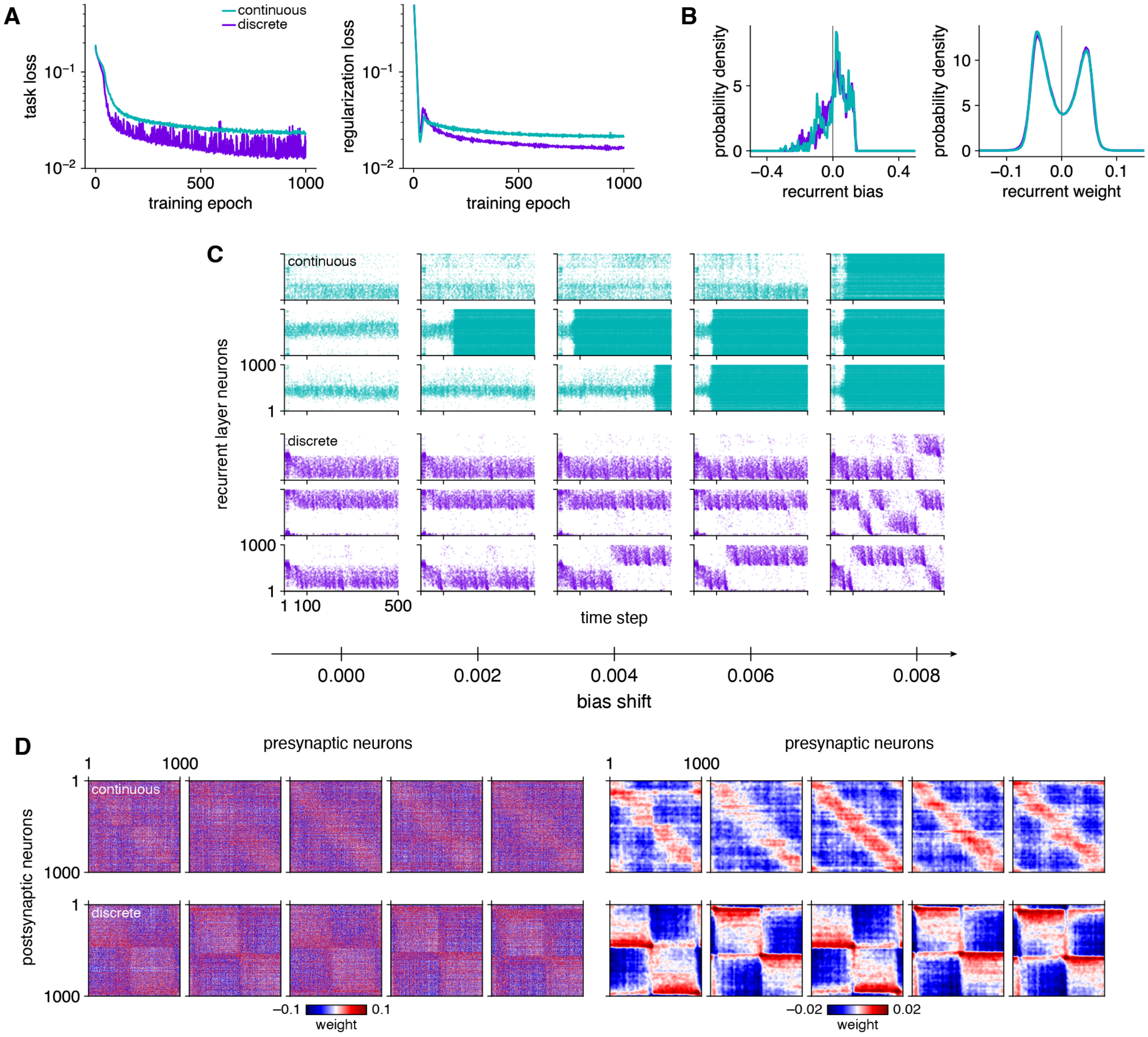


**Supplementary Figure 3**: Extended results for the continuous- and discrete-trained RNNs in Fig. 3 of the main text. (**A**) Task loss and total regularization loss during training. (**B**) Distributions of recurrent layer biases and weights after training. (**C**) Recurrent layer spike rasters over a range of bias shifts. Each row corresponds to one of three replicate continuous-trained or three replicate discrete-trained RNNs. (**D**) Left, sorted weight matrices for five replicate continuous-trained and five replicate discrete-trained RNNs. Right, weight matrices on the left smoothed with a 2D Gaussian filter of width 10 neurons with periodic boundaries.

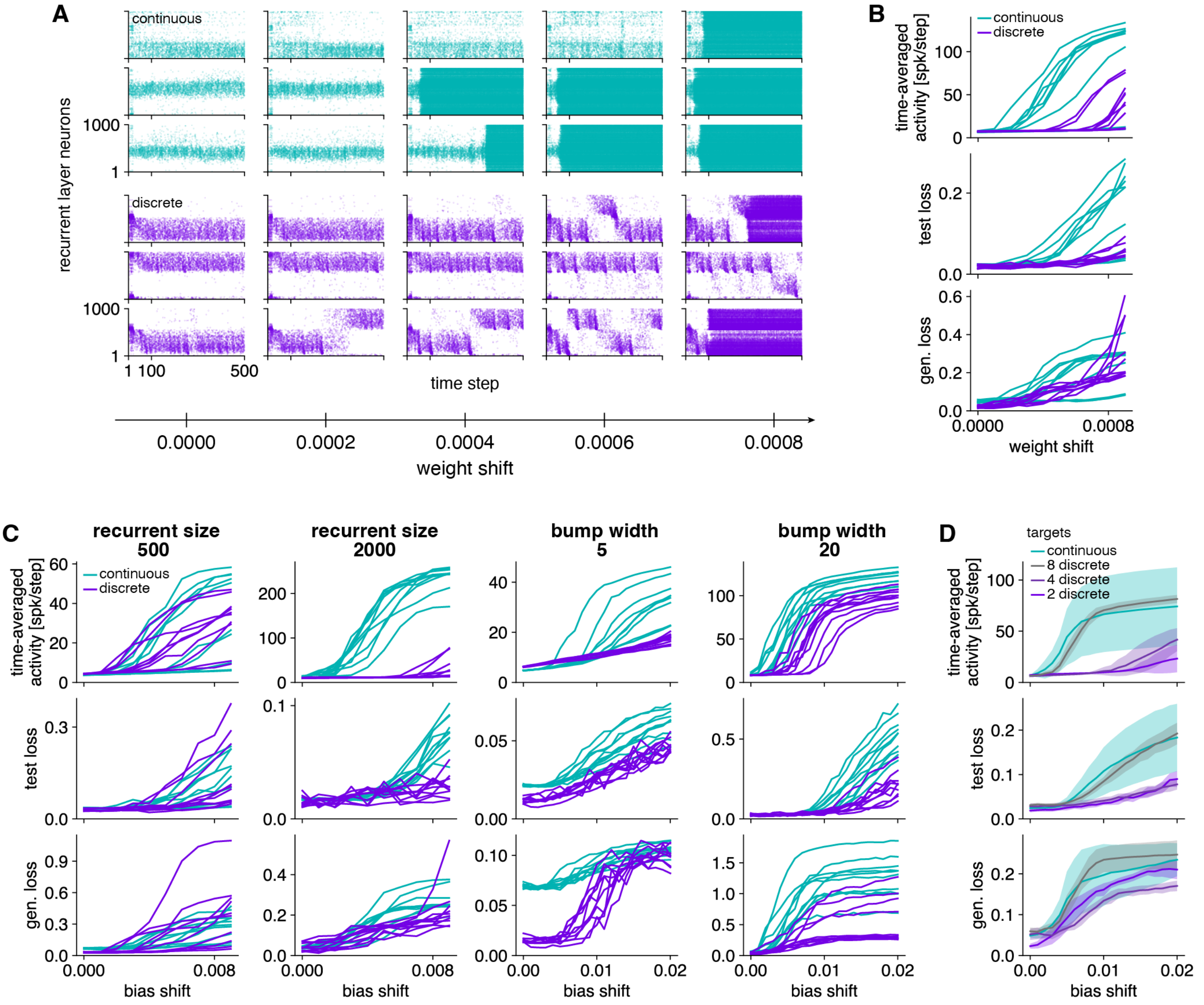


**Supplementary Figure 4**: Continuous- and discrete-trained RNNs with different perturbations and hyperparameters. (**A**,**B**) Perturbing networks by shifting all recurrent weights. (**A**) Recurrent layer spike rasters over a range of weight shifts. Each row corresponds to one of three replicate continuous-trained or three replicate discrete-trained RNNs. (**B**) Top, time-averaged population activity; middle, test loss; and bottom, temporal generalization loss. Data from 10 replicate simulations, and each line corresponds to one replica. (**C**) Similar to **B**, but for bias shift and the following changes in parameter values compared to those used in Fig. 3 of the main text: smaller and larger recurrent layer widths compared to 1000, and smaller and larger input and target bump widths compared to 10. (**D**) Behavior of discrete-trained RNNs approaches that of continuous-trained RNNs as the number of targets increases. Similar to **B**, but for different numbers of targets for the discrete task, with results from the continuous task shown for comparison. Lines and shaded areas indicate means and standard deviations over 10 replicate simulations.

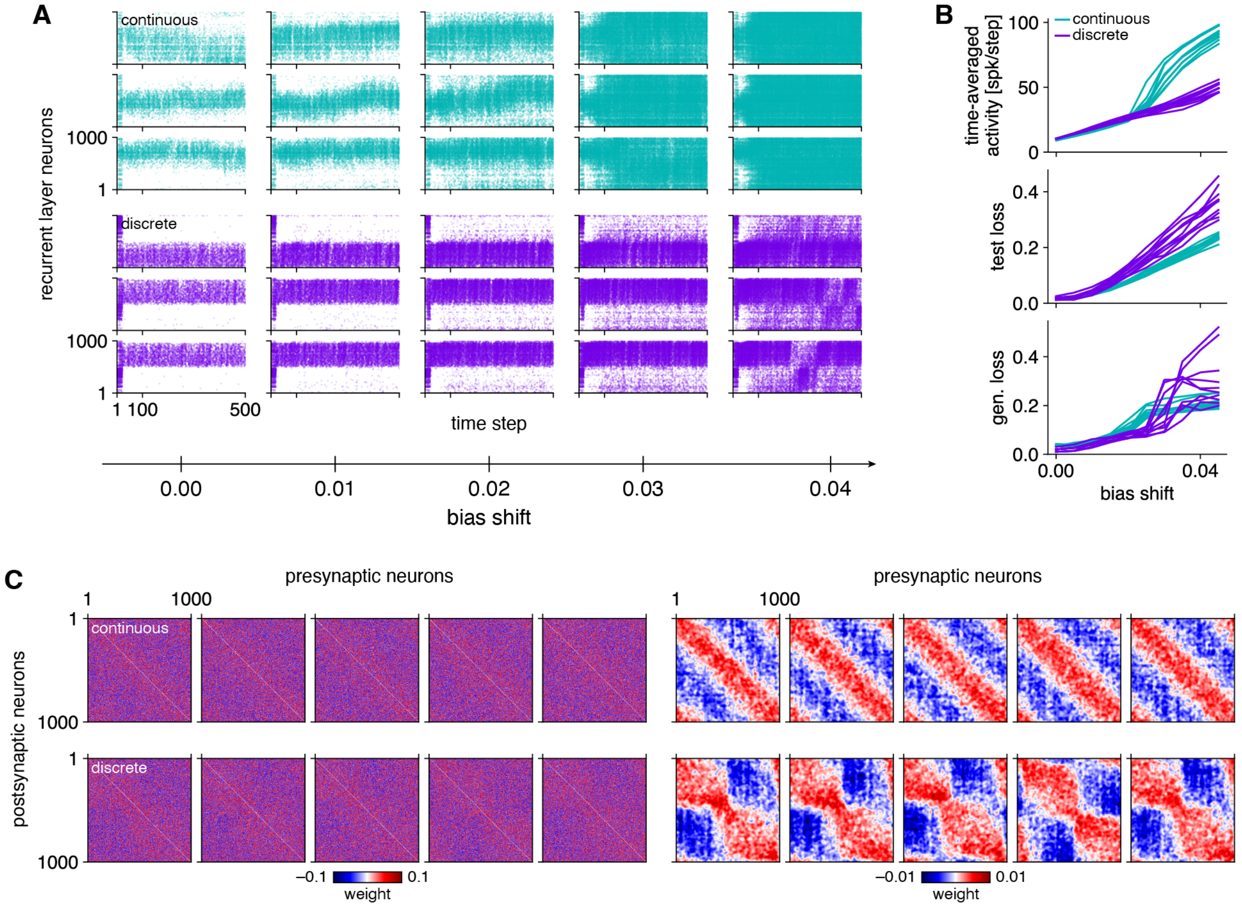


**Supplementary Figure 5**: Continuous- and discrete-trained RNNs with a different population activity regularizer. This regularizer encourages a target population activity at every time step instead of a target time-averaged population activity, which was used in Fig. 3 of the main text. (**A**,**B**) Perturbing networks by shifting all recurrent biases. (**A**) Recurrent layer spike rasters over a range of bias shifts. Each row corresponds to one of three replicate continuous-trained or three replicate discrete-trained RNNs. (**B**) Top, time-averaged population activity; middle, test loss; and bottom, temporal generalization loss. Data from 10 replicate simulations, and each line corresponds to one replica. (**C**) Left, sorted weight matrices for five replicate continuous-trained and five replicate discrete-trained RNNs. Right, weight matrices on the left smoothed with a 2D Gaussian filter of width 10 neurons with periodic boundaries.

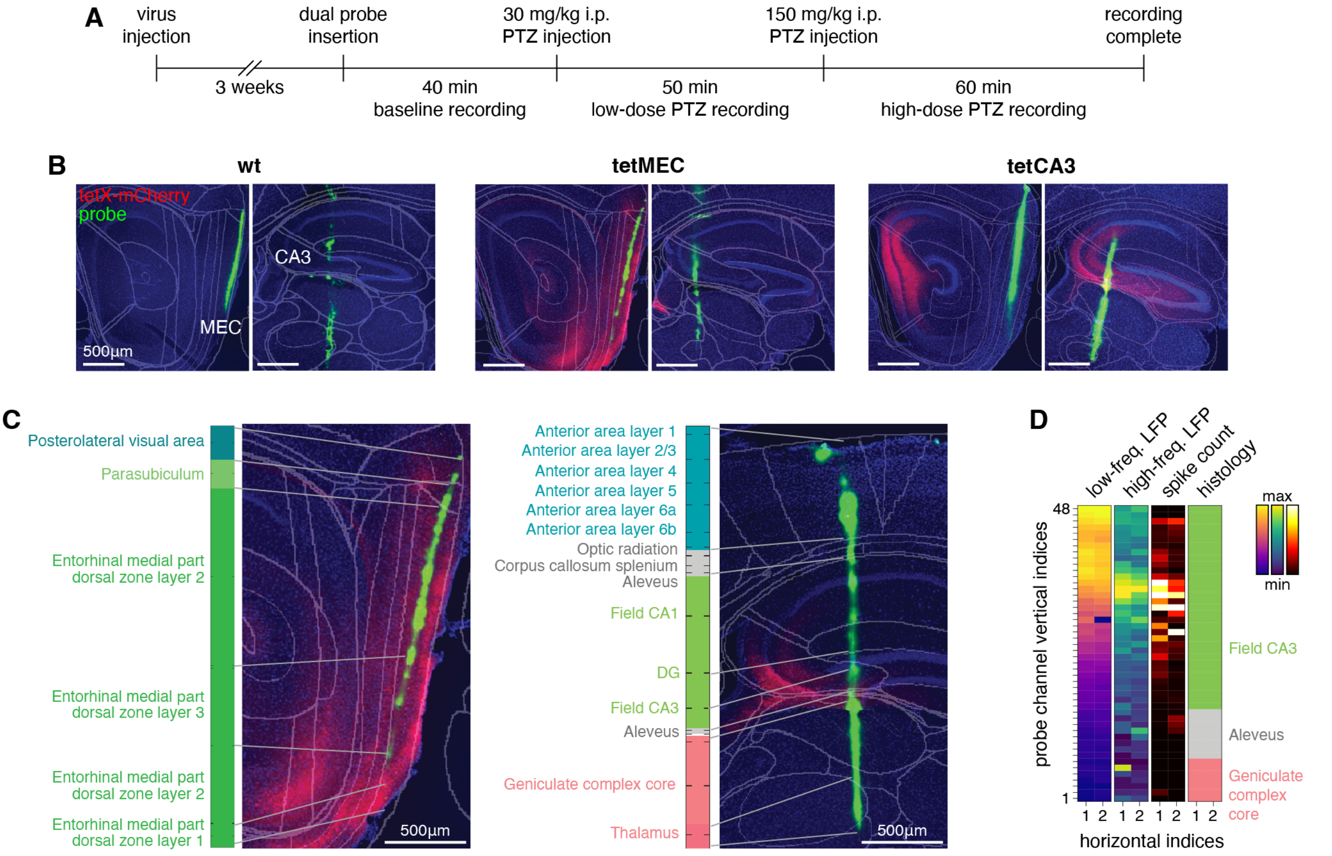


**Supplementary Figure 6**: Histological registration of brain anatomy and probe trajectory. (**A**) Schematic of the experimental protocol. (**B**) Example sagittal sections stained with DAPI (blue) illustrating the tracks of DiO-coated probes (green) and brain regions expressing the tetX-mCherry viral construct (red). Registration to the Allen Mouse Brain Atlas is overlaid (gray outlines). (**C**) Example mappings from probe trajectories to brain regions. Fluorophore colors are the same as those in **B**. (**D**) Example alignment of electrophysiological and histological properties used for probe channel selection. From left to right, columns indicate each channel's root-mean-square LFP amplitude between 1–500 Hz, root-mean-square LFP amplitude between 0.3–30 kHz, spike count, and histological label. Data from one shank of a Neuropixels 2.0 probe.

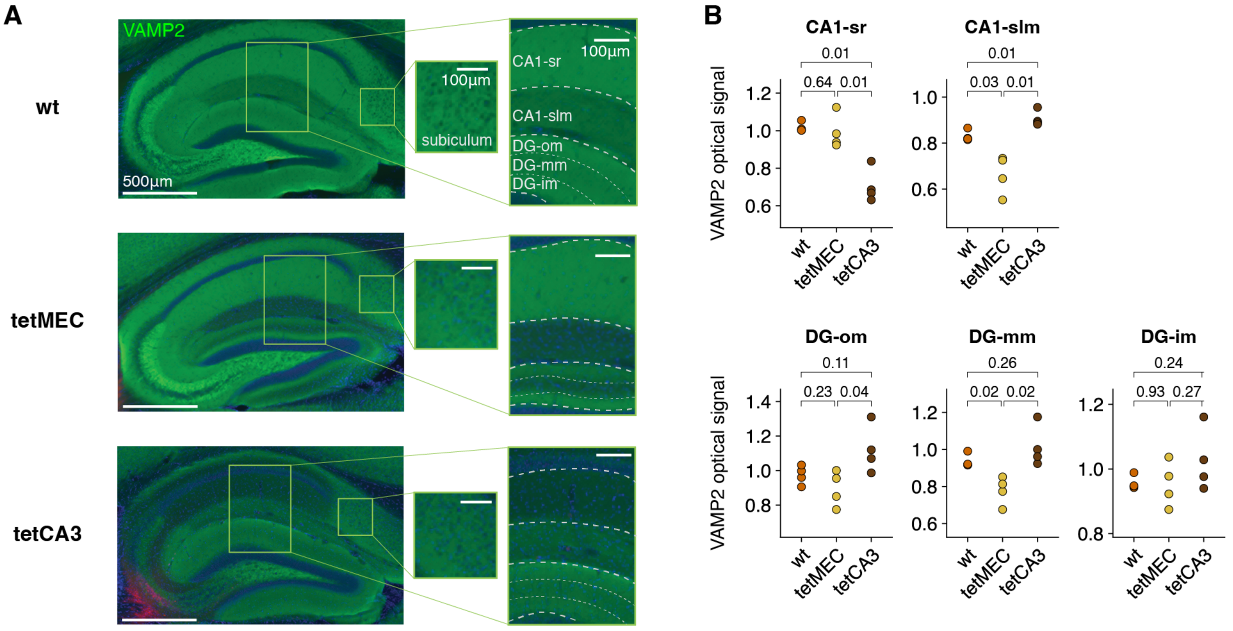


**Supplementary Figure 7**: Layer-specific VAMP2 cleavage by expressed tetanus toxin. (**A**) Example sagittal sections after immunohistochemical staining with anti-VAMP2 primary and Alexa Fluor 488 secondary antibodies (green). (**B**) tetMEC mice exhibit decreased VAMP2 optical signal in CA1-slm and DG-mm, which contain the axon terminals of MEC projections. tetCA3 mice exhibit decreased VAMP2 optical signal in CA1-sr, which contains the axon terminals of CA3 projections. The signal in each region is normalized by that of the subiculum. Each point corresponds to one animal. Data are from 4 wt, 4 tetMEC, and 4 tetCA3 mice. Numbers above brackets indicate $p$-values for unpaired two-sample $t$-tests comparing means. slm: stratum lacunosum-moleculare, sr: stratum radiatum, DG: dentate gyrus, om: outer molecular layer, mm: middle molecular layer, im: inner molecular layer.

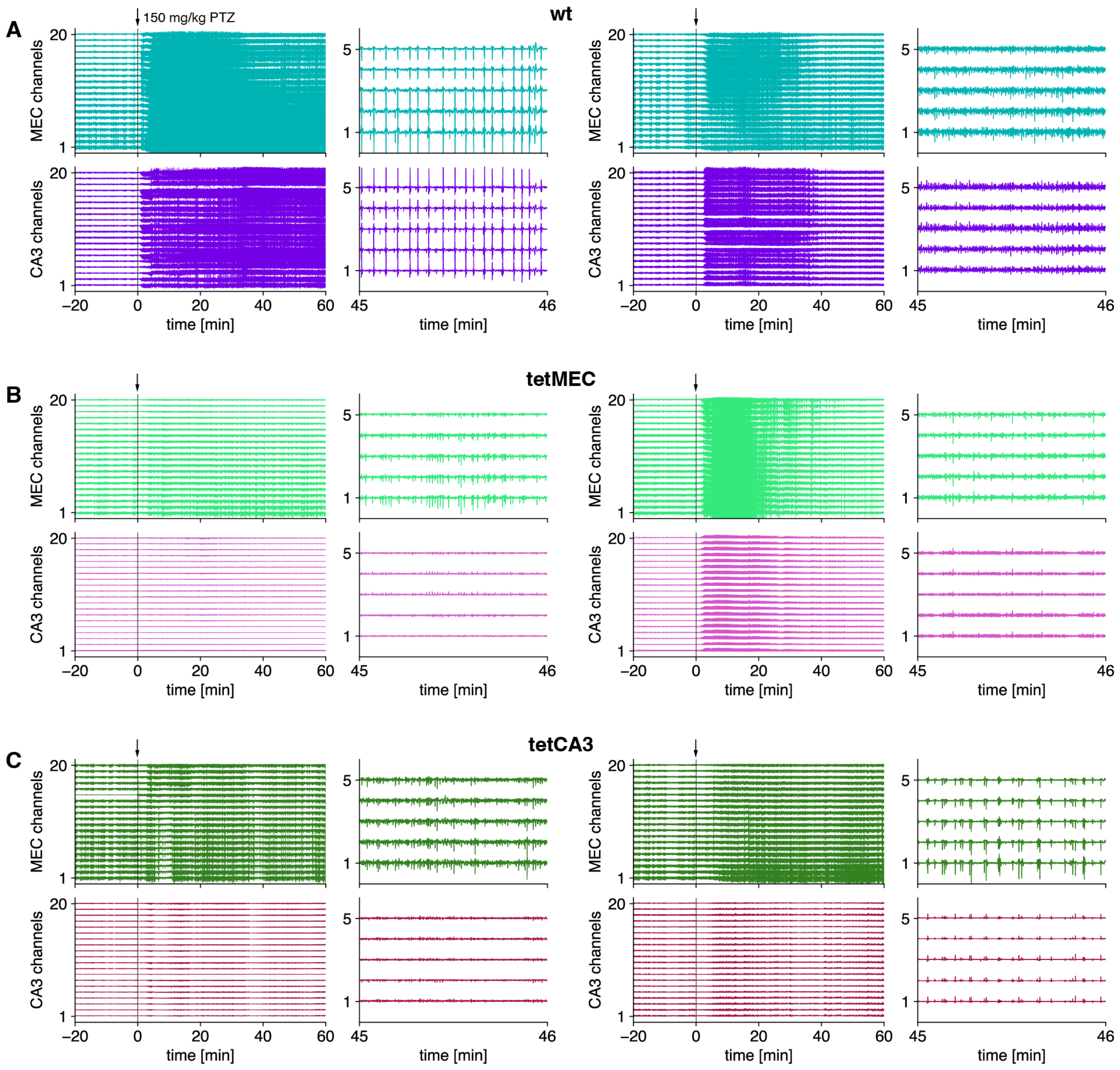


**Supplementary Figure 8**: Additional examples of MEC and CA3 LFP signals under high-dose PTZ. (**A**) LFP signals from evenly sampled channels in MEC and CA3 in 2 wt mice. A high-dose injection of PTZ is given at 0 min. (**B**) Similar to **A**, but for 2 tetMEC mice. (**C**) Similar to **A**, but for 2 tetCA3 mice.

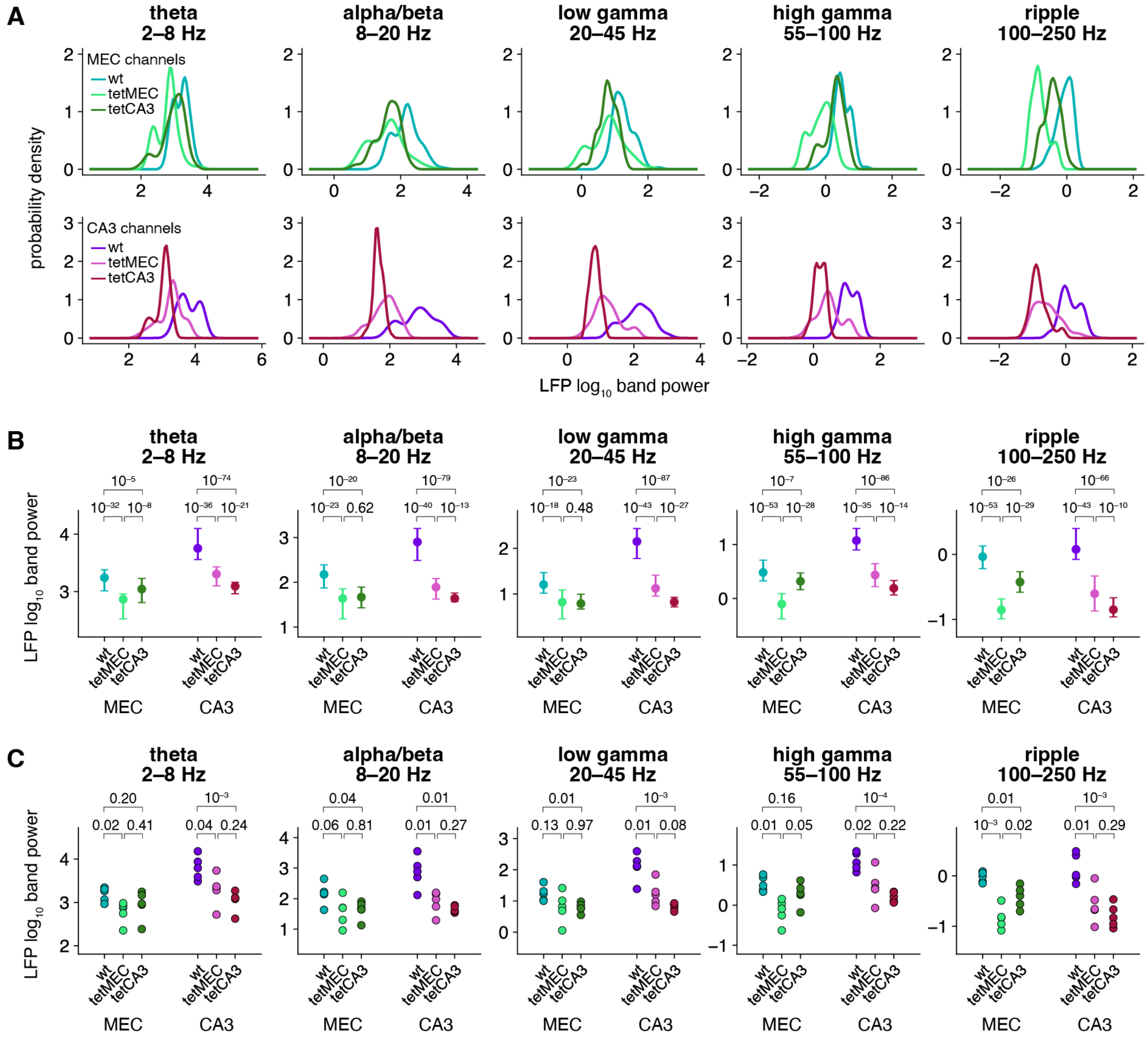


**Supplementary Figure 9**: MEC and CA3 LFP power within frequency bands under high-dose PTZ. (**A**) Kernel densities of LFP band power per channel 30–60 min after PTZ injection, aggregated across animals. (**B**) Medians and interquartile ranges of the data in **A**. (**C**) Medians of LFP band power per animal. **A**–**C** show data from 5 wt, 5 tetMEC, and 5 tetCA3 mice. In **B**, numbers above brackets indicate $p$-values for Mood's tests comparing medians. In **C**, numbers above brackets indicate $p$-values for unpaired two-sample $t$-tests comparing means.

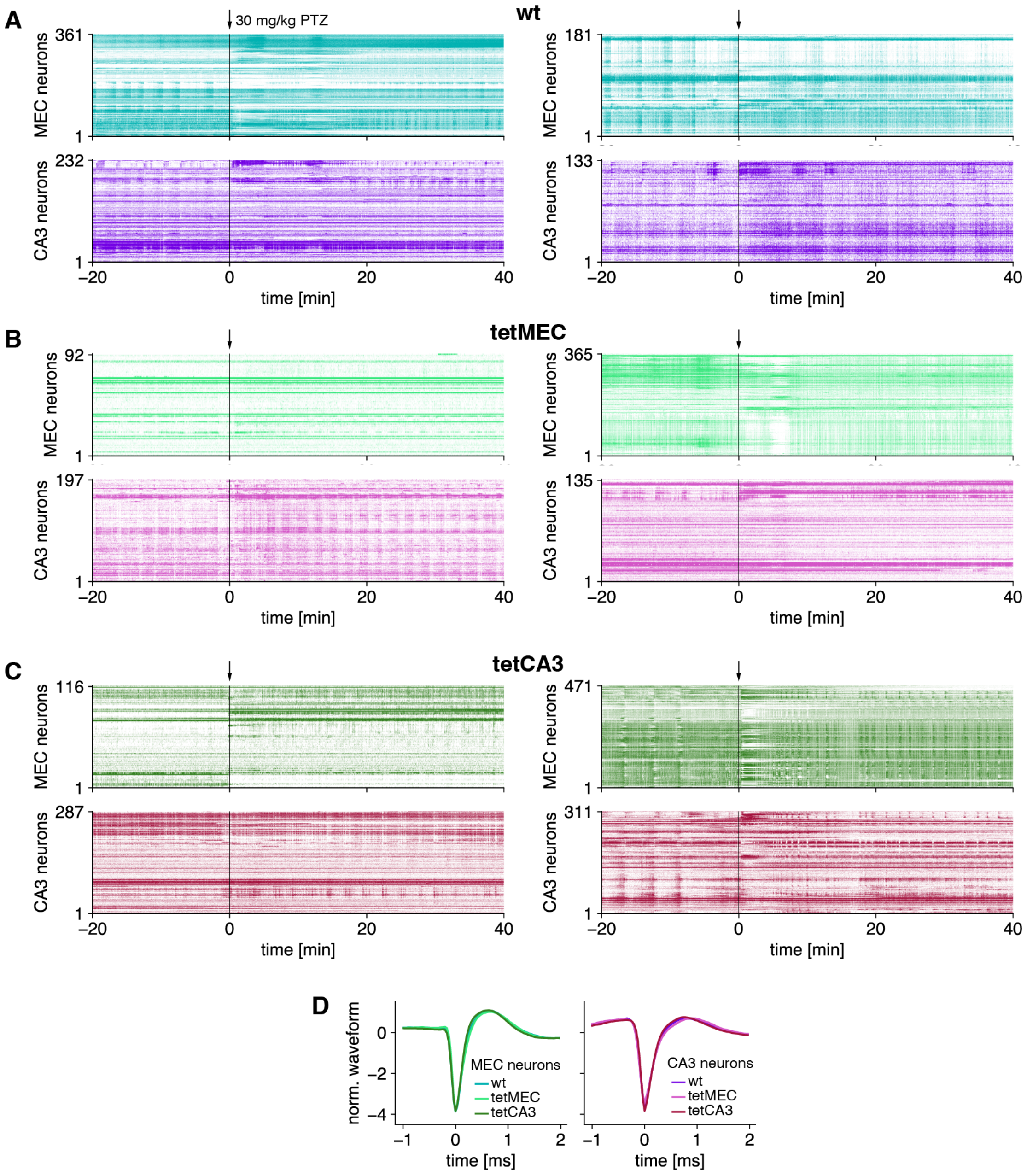


**Supplementary Figure 10**: Additional examples of MEC and CA3 spike rasters under low-dose PTZ. (**A**) Single-unit spike rasters for MEC and CA3 in 2 wt mice. A low-dose injection of PTZ is given at 0 min. (**B**) Similar to **A**, but for 2 tetMEC mice. (**C**) Similar to **A**, but for 2 tetCA3 mice. (**D**) Spike waveform templates divided by their standard deviation, averaged across neurons and animals. The channel with largest trough-to-peak amplitude is chosen for each template, and troughs are aligned to 0 ms. Data are from 6 wt, 6 tetMEC, and 6 tetCA3 mice.

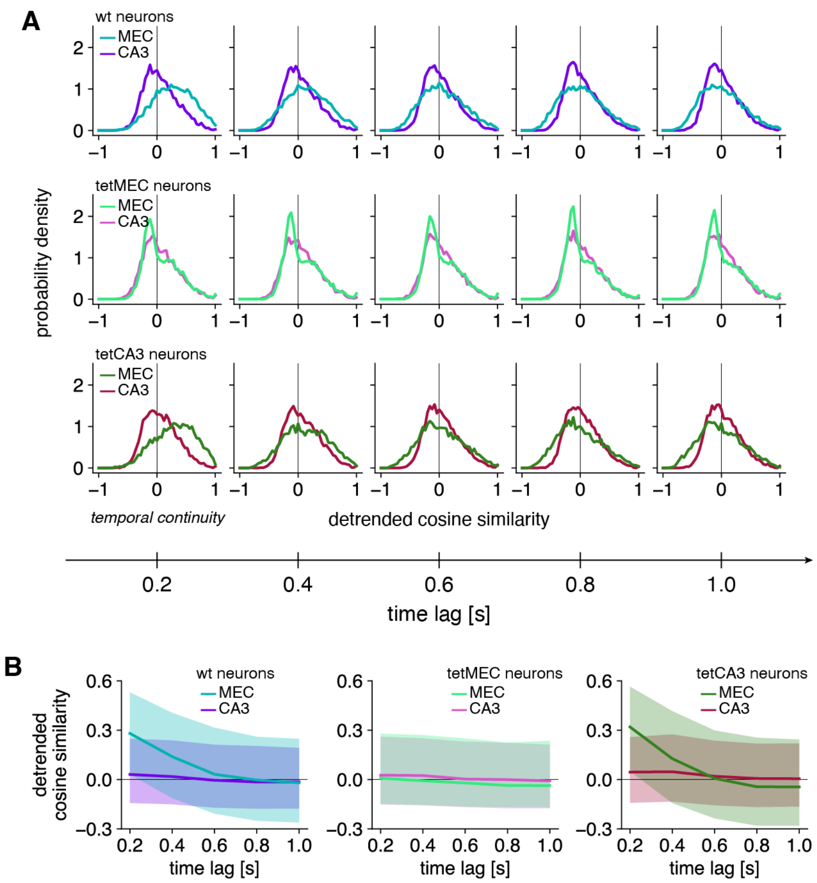


**Supplementary Figure 11**: Extent of temporal overlap in MEC and CA3 network states under low-dose PTZ. (**A**) Histograms of cosine similarity between detrended network states separated by various time lags. Time lag 0.2 s corresponds to our definition of temporal continuity. (**B**) Medians (lines) and interquartile ranges (shaded areas) of the data in **A**. **A**, **B** show data from 6 wt, 6 tetMEC, and 6 tetCA3 mice.
